# Engineering filamentous myosins for optical control of contractility

**DOI:** 10.1101/2025.08.19.666446

**Authors:** Sasha Zemsky, Paul V. Ruijgrok, Zev Bryant

**Affiliations:** Department of Bioengineering, Stanford University, Stanford, CA, USA; Graduate Program in Biophysics, Stanford University, Stanford, CA, USA; Bio-X Institute, Stanford University, Stanford, CA, USA; Department of Structural Biology, Stanford University, Stanford, CA, USA

## Abstract

Understanding the behaviors of contractile actomyosin systems requires precise spatiotemporal control of filamentous myosin activity. Here, we develop a tool for optical control of contractility by extending the MyLOV family of gearshifting motors to create engineered filamentous myosins that change velocity in response to blue light. We characterize these minifilaments using *in vitro* single-molecule tracking assays, contractility assays in reconstituted actin networks, and imaging of contractile phenotypes in *Drosophila* S2 cells. The minifilaments change speed and/or direction when illuminated, display speeds that fall within and beyond the relevant physiological range, and display high processivities. Additionally, minifilament-driven contraction rates increase in blue light both *in vitro* and in S2 cells. Finally, we develop an alternative design for minifilaments that only interact processively with actin in blue light. Engineered minifilaments can be used to dissect behaviors such as self-organization and mechanotransduction in contractile systems both *in vitro* and in cells and tissues.

## Introduction

Nonmuscle myosin II (NMII) filaments generate forces and remodel the actin cytoskeleton in eukaryotic cells to drive processes such as cell migration, cell adhesion, cytokinesis, and tissue morphogenesis^1–3^. Manipulation of NMII activity both *in vivo* and *in vitro* has been used to dissect the role of NMII in these dynamic processes. *In vivo*, strategies such as gene deletions/mutations, RNA interference, and drug treatments have been used to directly target NMII or to target signaling proteins that modulate NMII activity^1,2^. *In vitro*, networks or bundles of actin and myosin II have been used as minimal models of contractile systems. In these reconstitutions, the concentration of myosin II, the minifilament size, the type of myosin II isoform, and the concentration of actin crosslinking proteins can be altered to investigate how contractility occurs and how it is organized and regulated in actomyosin systems without sarcomeric organization^4–9^. However, understanding how specific patterns of NMII contractility in space and time give rise to cytoskeletal reorganization, force exertion, and mechanotransduction requires techniques that enable control of contractility with high spatiotemporal precision^10,11^ both in living cells and in reconstituted systems.

Multiple methods have been developed to control contractility in cells and tissues with high spatiotemporal precision using optogenetics. Optogenetic modules have been fused to several different proteins that participate in signaling pathways that regulate contractility^10,12–14^. The most common target is RhoA, a signaling GTPase that promotes contractility by reorganizing actin and phosphorylating the NMII regulatory light chain (RLC). Optical control of RhoA has been used to manipulate cytokinesis^15^, cell-cell adhesion^16^, exertion of traction forces^17,18^, cell migration^19^, and morphogenesis^20^. However, these optogenetic tools do not specifically target NMII, as RhoA has multiple downstream effectors^21^. Additionally, these methods have never been applied to *in vitro* active matter systems — and it would be challenging to reconstitute signaling pathway components and achieve reversible spatiotemporal control over downstream NMII activity *in vitro*.

Optical control of contractility *in vitro* has been explored previously using light-induced activation of photocaged calcium^22^ and light-induced inactivation of the myosin II inhibitor blebbistatin^23–25^. However, in these systems the optical activation is irreversible at the molecular scale. These methods also have limited applicability in cells. Blebbistatin photoinactivation produces free radicals during extended blue light exposure, and active blebbistatin can diffuse into the cell from the surrounding solution once the blue illumination has stopped^26^. Thus, each of the methods for optical control of NMII described here has limitations either *in vitro* or in cells. An ideal method would be applicable both in cells and *in vitro* so that experimental results could be compared across both types of systems.

An alternative strategy is to place a module for optical control directly into the myosin protein rather than targeting upstream signaling proteins. This was achieved previously by engineering a gearshifting lever arm that changes geometry in response to blue light, leading to a change in myosin velocity *in vitro*^27,28^. Gearshifting myosin XI tetramers were used for temporal control of cargo transport in cells^28^ and for spatiotemporal control of the dynamics of an extensile actin liquid crystal^29^. However, light-activated gearshifting has not yet been applied to filamentous myosins, and it has not been used to control contractility. Applying this strategy to filamentous myosins would provide a tool to specifically activate myosin contractility without affecting other protein functions. This tool could also be used in minimal reconstituted systems, which would provide a unique opportunity to use the same tool to manipulate contractility both *in vitro* and in cells. Finally, the biophysical parameters of engineered myosin filaments could be tuned by modifying the motor and/or tail domains or the optical dose^28^, which could enable more detailed exploration of how myosin properties influence contractile behaviors.

Here, we apply light-activated gearshifting to filamentous myosins. We build hybrid myosin filaments using motor domains from myosin V, VI, or XI and tail domains from NMIIB or *zipper.* We use *in vitro* motility assays to demonstrate optical modulation of minifilament velocities, and we use a minimal *in vitro* contractility assay to demonstrate optical control of contraction rates. In addition, we develop an alternative design strategy that uses light-induced dimerization to control minifilament assembly, and we show that these constructs can also be used for optical control of contractility *in vitro.* Finally, we show that engineered minifilaments can be used to control contraction of protrusions in live *Drosophila* S2 cells.

## Results

### Design of gearshifting minifilament constructs

The NMII heavy chain consists of a motor domain, a lever arm, and a tail domain. Multiple NMII molecules assemble into bipolar minifilaments via interactions between their tail domains^1–3^. Isoforms of NMII found in animals typically contain ∼30 to ∼60 motor domains per filament^30,31^ and move with velocities ranging from ∼40 nm/s for human NMIIB^32^ to ∼250 nm/s for *Drosophila* NMII^31^.

To design gear-shifting minifilaments, we replaced the native NMII lever arm with an engineered lever arm (described previously in ref. ^27,28^) consisting of a Light-Oxygen-Voltage 2 (LOV2) domain flanked by 3-helix bundle domains from alpha actinin and spectrin (Fig. 1a, Supp. Fig. 1). In the absence of blue light, the lever arm is designed to fold into a rigid hairpin structure. In the presence of blue light, the Jα helix of the LOV2 becomes disordered, and the portion of the lever arm C-terminal to the LOV2 no longer contributes to the lever arm swing, which leads to a change in the size and/or direction of the power stroke (Fig. 1b). This effect is demonstrated by structural models of myosin VI and myosin XI motor domains fused to the engineered lever arm (Supp. Fig. 2), which provide idealized predictions for the stroke vectors in the dark and lit states. For each construct, the predicted stroke vector in the lit state has a similar magnitude but completely reversed direction in comparison to the stroke vector in the dark state. The models therefore suggest that in response to blue light, both constructs would maintain approximately the same speed but would completely switch their direction of motion along actin. However, previous experimental results^27,28^ demonstrate that some constructs display reduced speed rather than reversed direction in the dark state, which suggests that the light-induced velocity modulation can be smaller than predicted by the model for certain constructs. This quantitative difference between modeling and experimental results may be due to compliance in the lever arm, and/or differences between the exact geometry of the lever arm and the geometry predicted by the model.

**Figure 1.**
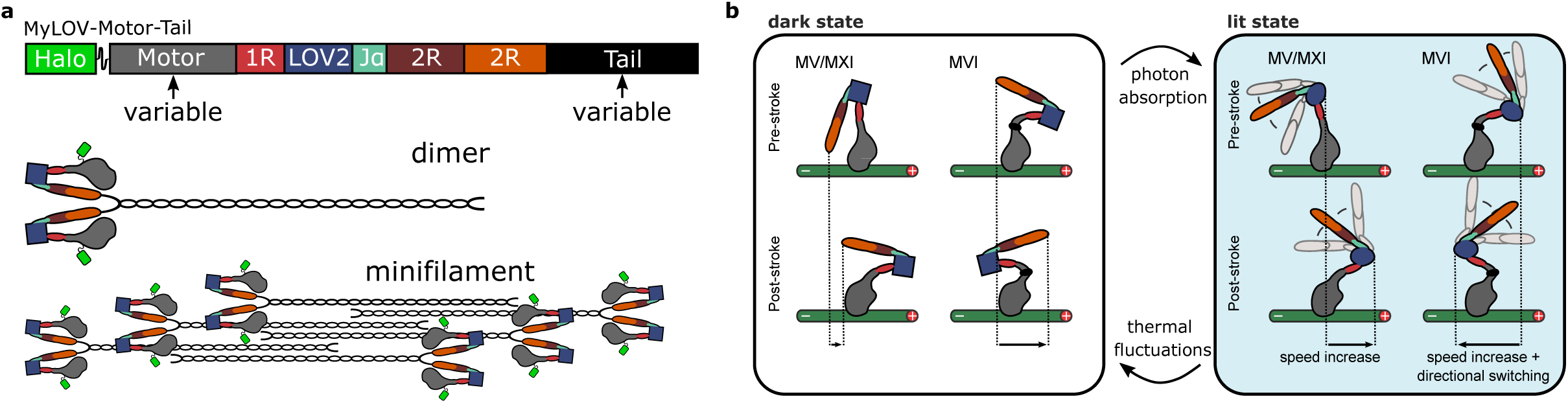
Optical control of minifilament velocity *in vitro*. **a**) Block diagram of an optically controllable gearshifting minifilament construct (top) and cartoon of a dimer (center) and dimers assembled into a minifilament (bottom). **b**) Diagram of the proposed mechanism for optically-controllable velocity switching.

The gearshifting design is modular, as the engineered lever arm can be fused to different motor domains^27^ and different multimerization domains^28^. Previous light-activated gearshifting designs contained motor domains from myosin VI and myosin XI^27^ and were either monomeric or tetrameric^28^. To form minifilaments, we fused the engineered lever arm to the full-length tail domain from human NMIIB starting at L844 (Fig. 1a, Supp. Fig. 1), which is immediately after the invariant proline that marks the beginning of the tail domain^33^. We replaced the NMIIB motor domain with motor domains from myosin VI, *Nicotiana tabacum* myosin XI, and myosin Va. These constructs are abbreviated as MyLOV-VI-IIB, MyLOV-XI-IIB, and MyLOV-V-IIB. To achieve a greater range of velocities, we introduced mutations into the myosin VI and myosin XI motor domains. MyLOV-VI_L310G_-IIB contains an L310G mutation at insert-1 in the myosin VI nucleotide binding site, which increases myosin VI velocity and reduces processivity^34^. MyLOV-XI_L2+4_-IIB has 4 lysines inserted into loop 2 of the myosin XI motor domain, which was previously shown to reduce the velocity of *Chara corallina* myosin XI by more than a factor of 4^35^. Finally, to determine whether the identity of the NMII tail domain influences the behavior of the engineered minifilaments, we replaced the NMIIB tail in MyLOV-VI-IIB with the tail from *Drosophila* NMII (*zipper*) to create the construct MyLOV-VI-*zip*.

### Processive single-molecule motility assays on individual actin filaments

We measured the velocity and optical modulation of the minifilament constructs using an *in vitro* processive single-molecule motility assay that was used previously to characterize the processive motion of wildtype NMII minifilaments^32,36^. In this assay, minifilaments walk along biotinylated actin filaments that are attached to a streptavidin-coated coverslip (Fig. 2a). Motor velocities were measured using kymograph analysis (see Methods). To determine motor directionality, we identified the plus and minus-end of each actin filament by imaging a previously-characterized myosin VI tetramer (M6DI_816_2R∼TET) that always walks toward the minus-end of actin^37^.

**Figure 2.**
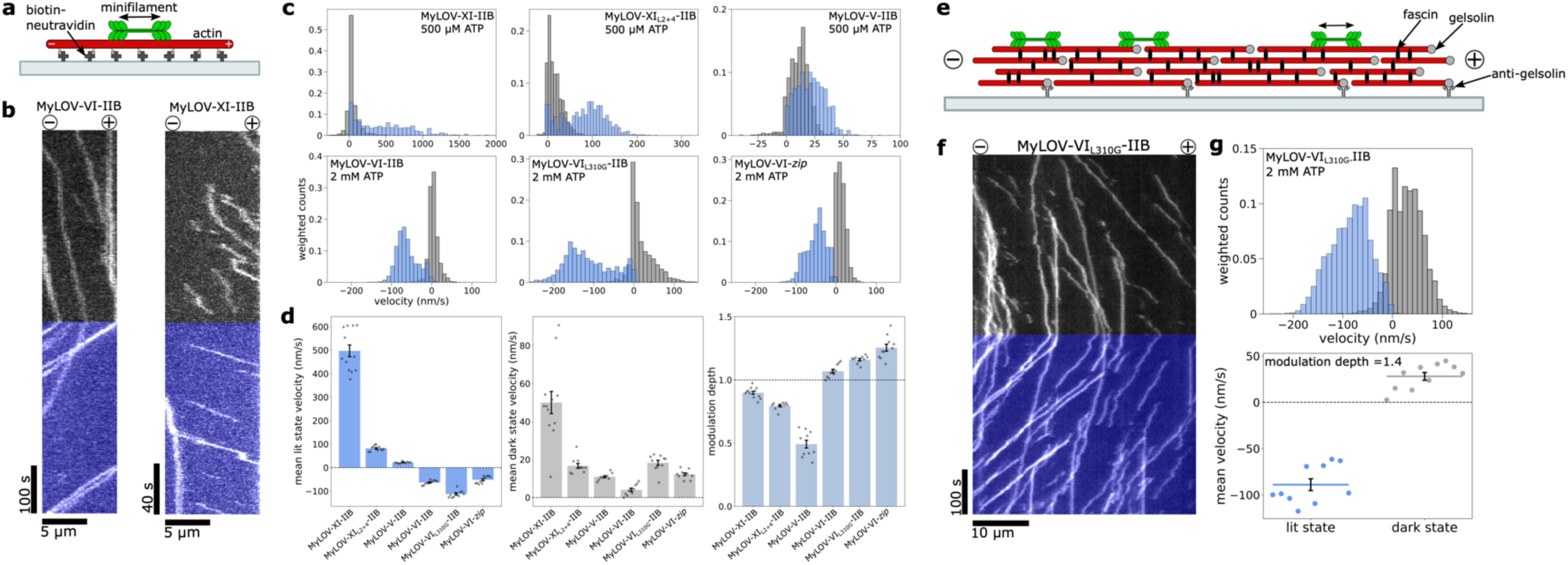
Velocities of gearshifting minifilaments in processive motility assays. **a**) Schematic of *in vitro* processive motility assay on single biotinylated actin filaments. **b)** Example kymographs on a biotinylated actin filament for MyLOV-VI-IIB at 2 mM ATP (left, see also Supp. Movie 1) and MyLOV-XI-IIB at 100 μM ATP (right). The right sides of the kymographs represent the plus-ends of the actin, and the left sides represent the minus-ends. Blue regions indicate periods of blue illumination. **c**) Normalized time-weighted histograms of velocities of individual runs in processive motility assays for each of the 6 constructs described in the text. Construct name and ATP concentration are labeled on the histograms. Gray bars represent runs that occurred in the dark and blue bars represent runs with blue illumination. Negative velocities are minus-end-directed and positive velocities are plus-end-directed. Runs with velocities above 2000 nm/s for MyLOV-XI-IIB (∼3% of all lit state runs) and below –260 nm/s for MyLOV-VI_L310G_-IIB (∼0.05% of all lit state runs) were cut off in the histograms for data visualization purposes. **d**) Average time-weighted lit state velocity (left), time-weighted dark state velocity (middle), and modulation depth (defined as 1 – v_dark_/v_lit_, bottom) for all constructs. Error bars show standard error. Each gray point represents an average across all runs from one experiment. Final averages and standard errors were calculated across experiments. **e**) Schematic of *in vitro* processive motility assay on aligned and polarized fascin-actin bundles. **f)** Example kymograph on a fascin-actin bundle for MyLOV-VI_L310G_-IIB (see also Supp. Movie 4). **g)** Normalized time-weighted histogram of velocities of individual runs (top) and average time-weighted lit and dark state velocities calculated across all experiments (bottom) for MyLOV-VI_L310G_-IIB on fascin-actin bundles. Each point in the bottom plot represents an average across all runs from one experiment, and error bars show standard error. All data shown in **c**, **d**, and **g** was collected from 10 to 12 separate experiments over at least 2 days with 2 protein preps for each construct. See Supp. Table 1 for the total number of runs measured for each construct.

All of the gearshifting minifilament constructs change velocity in response to blue light (Fig. 2b-d, Supp. Table 1). The relative change between the dark state velocity (v_dark_) and the lit state velocity (v_lit_) for each construct can be represented by the modulation depth^28^, defined as 1 – v_dark_/v_lit_ (Fig. 2d). The velocity and run length results for all constructs are summarized in Supp. Table 1.

MyLOV-XI-IIB, MyLOV-XI_L2+4_-IIB, and MyLOV-V-IIB are primarily plus-end-directed both in the dark and lit states (Fig. 2c). MyLOV-V-IIB has a two-fold increase in velocity in the presence of blue light (v_dark_ = 11 nm/s, v_lit_ = 22 nm/s). MyLOV-XI-IIB is faster and has a greater fold-change in velocity, switching from v_dark_ = 50 nm/s to v_lit_ = 497 nm/s (Supp. Movie 1). MyLOV-XI_L2+4_-IIB is slower than MyLOV-XI-IIB in both states (v_dark_ = 17 nm/s, v_lit_ = 82 nm/s). The ATP concentration used in these assays (500 μM) was slightly below the saturating ATP concentration for MyLOV-XI-IIB. However, the velocity at saturating ATP can be extrapolated by using a Michaelis-Menten fit to velocity vs. [ATP] data that was collected previously for an engineered construct with the same motor domain^37^. At saturating ATP, the predicted velocities for MyLOV-XI-IIB are v_dark_ = 60 nm/s and v_lit_ = 600 nm/s. Thus, MyLOV-XI-IIB in the lit state is ∼4-5 times faster than the fastest human NMII isoform^32^ and ∼2 times faster than *Drosophila* NMII^31^.

MyLOV-VI-IIB, MyLOV-VI_L310G_-IIB and MyLOV-VI-*zip* all switch from plus-end-directed to minus-end-directed in response to blue light (Fig. 2c). MyLOV-VI-IIB switches from v_dark_ = 4 nm/s to v_lit_ = –63 nm/s (Supp. Movie 2). MyLOV-VI_L310G_-IIB is faster than MyLOV-VI-IIB in both states (v_dark_ = 18 nm/s, v_lit_ = –112 nm/s). MyLOV-VI-*zip* behaves similarly to MyLOV-VI-IIB (v_dark_ = 12 nm/s, v_lit_ = –52 nm/s, Supp. Movie 3). To our knowledge, these constructs are the first examples of minifilaments that can move toward the minus end of actin. In addition, in the lit state these constructs are more than an order of magnitude faster than previous optically controllable directional switching myosin VI constructs^27^.

All of the constructs walk processively along actin at saturating or near saturating ATP concentrations, with run lengths frequently reaching several microns (Supp. Fig. 3 and Supp. Table 1). MyLOV-XI_L2+4_-IIB, MyLOV-V-IIB, MyLOV-VI-IIB and MyLOV-VI-*zip* were rarely observed to dissociate from actin (Supp. Table 1) and therefore we did not calculate average run lengths for these constructs. MyLOV-VI_L310G_-IIB dissociates from actin more frequently than MyLOV-VI-IIB but still has an average run length, L, of several microns in both the dark and lit states (L_dark_ = 7.15 μm, L_lit_ = 22.29 μm). MyLOV-XI-IIB is the least processive of the 6 constructs (L_dark_ = 0.73 μm, L_lit_ = 2.21 μm). Overall, we have shown that the engineered minifilaments can walk processively on actin and can change speed and/or direction in response to blue light.

### Processive single-molecule motility assays on fascin-actin bundles

In addition to assays on individual actin filaments, we also performed motility assays on bundles of actin and fascin, which crosslinks actin filaments in a parallel orientation to form unipolar bundles^38^. Recently, it was shown that NMII filaments can move processively on parallel actin bundles in cells^39^. A minimal *in vitro* reconstitution of parallel actin bundles could improve our understanding of how minifilaments behave on these actin geometries in cells.

We used a modified version of a previously-described protocol^40^ to produce bundles of fascin and actin that were tightly attached to the surface and also aligned and polarized along the direction of flow, with plus-ends oriented toward the source of the flow/the inlet of the channel (Fig. 2e, Supp. Fig. 4a). This actin geometry is convenient for visualizing motor directionality, as plus-end-directed motors all move in one direction on the bundles (toward the inlet of the channel), and minus-end-directed motors all move in the opposite direction (toward the outlet of the channel), as demonstrated in Supp. Movie 4. We tested MyLOV-VI_L310G_-IIB in this assay and found that it can walk processively along bundles (Fig. 2f). It also switches directionality in response to blue light, with velocities similar to those measured on individual actin filaments (v_dark_ = 28 nm/s, v_lit_ = –89 nm/s, Fig. 2g and Supp. Movie 4).

To confirm that the bundles are polarized with plus-ends oriented toward the source of the flow, we performed identical experiments using the engineered myosin VI tetramer^37^ M6DI_816_2R∼TET (Supp. Fig. 5a). M6DI_816_2R∼TET is always minus-end-directed, so runs that are measured as minus-end-directed indicate that a bundle is polarized in the expected orientation, while plus-end-directed runs indicate the reverse orientation. The mean velocity of M6DI_816_2R∼TET on fascin-actin bundles is similar to the velocity reported previously for this construct on single actin filaments^37^. Only 15 out of 1303 runs (∼1%) were scored as plus-end-directed, indicating that most bundles are polarized in the expected orientation and that the directionality is measured correctly for close to 99% of runs. This conclusion is also supported by particle image velocimetry analysis of the motion of the motors along the actin bundles (Supp. Fig. 5b,c). Thus, we have shown that engineered minifilaments move processively on fascin-actin bundles *in vitro*, and this assay can be used to measure the velocities of bidirectional motors.

### Contractile bundle assays

We next determined if the engineered minifilaments can contract actin structures *in vitro* and if the contraction rates can be modulated using blue light. Recently, a minimal model of contractile actin bundles was used to study binding of LIM domain proteins to actin bundles under tension^41^. We developed a similar assay using bundles of actin and ɑ-actinin, which crosslinks actin in both parallel and antiparallel orientations to form mixed-polarity bundles^42^. The bundles were nonspecifically attached to a coverslip in the presence of methylcellulose (Fig. 3a). Segments of some bundles were immobilized on the coverslip, which constrained the bundle network to a 2D layer close to the coverslip that could be imaged in TIRF. Other segments of bundles remained detached from the surface and could be deformed by the minifilaments.

**Figure 3.**
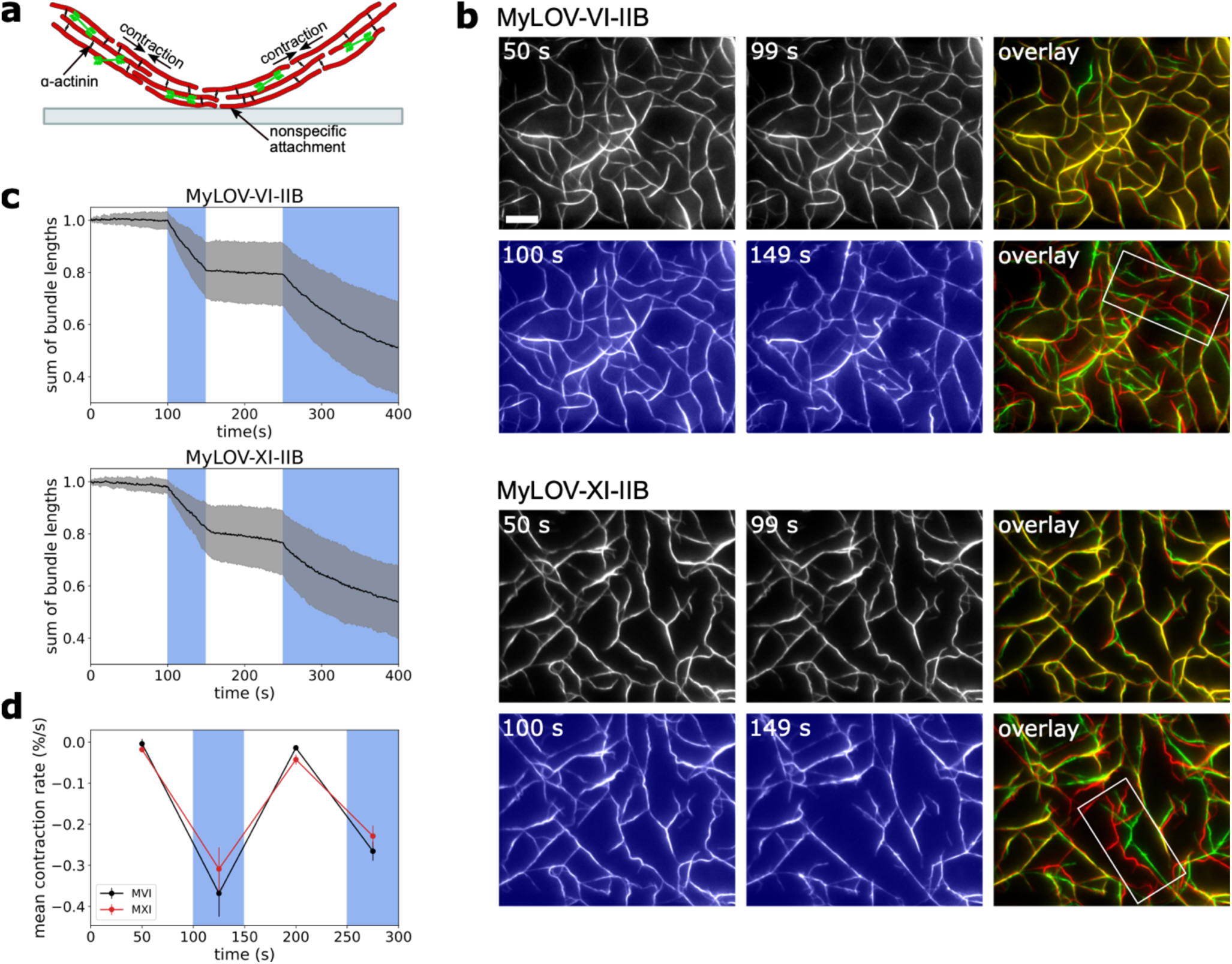
Optical control of contraction rate *in vitro*. **a**) Schematic of the *in vitro* contractility assay. **b**) Snapshots of actin bundles in contractility assays with MyLOV-VI-IIB (top, taken from Supp. Movie 6) and MyLOV-XI-IIB (bottom, taken from Supp. Movie 5) at 2 mM ATP showing the change in the actin bundle architecture during 50 s of dark (timepoints 50 s and 99 s) and 50 s of blue illumination (timepoints 100 s and 149 s). Overlays are shown for each 50 s time period with the earlier timepoint in red and the later timepoint in green. The scale bar represents 10 μm. White boxes in overlays highlight bundles that straighten during illumination. **c**) Sum of the lengths of all actin bundles over time in contractility assays with MyLOV-VI-IIB (top) and MyLOV-XI-IIB (bottom) at 2 mM ATP, normalized to the total bundle length in the first frame. Blue regions represent periods of blue illumination. The black curves are averages of 10 movies from 10 different experiments with 2 protein preps over multiple days. The gray envelopes represent ± one standard deviation. **d**) Mean contraction rates in the dark and light for MyLOV-VI-IIB (black) and MyLOV-XI-IIB (red) at 2 mM ATP. Contraction rates were calculated as the slopes of piecewise linear fits to the bundle length vs. time traces. Slopes were calculated for 4 time periods: times 0-99 s (dark), 100-149 s (lit), 150-249 s (dark), and 250-299 s (lit). Each point represents the mean slope (averaged over 10 movies) for each time period. Contraction rates are shown as the percentage change in total bundle length per second. Error bars represent standard errors.

MyLOV-VI-IIB and MyLOV-XI-IIB were tested in this assay. In each experiment, 2 periods of dark were alternated with 2 periods of blue illumination. Both constructs cause actin bundles to straighten and shorten in response to blue light (Fig. 3b and Supp. Movies 5 and 6). To calculate the bundle contraction rate, the sum of the lengths of all bundles in the field of view was measured for each frame of a movie (Fig. 3c and Supp. Fig. 7). The average normalized rate of change of the total bundle length, r (measured in percent change per second), was calculated for each period of dark and each period of blue illumination (Fig. 3d). For MyLOV-VI-IIB, r increases in magnitude from r_dark,1_ = –0.004 ± 0.012%/s (mean ± s.e.m) to r_lit,1_ = –0.368 ± 0.057%/s for the first cycle of dark and blue illumination, and from r_dark,2_ = –0.014 ± 0.005%/s to r_lit,2_ = –0.266 ± 0.023%/s for the second cycle. MyLOV-XI-IIB displays similar behavior (r_dark,1_ = – 0.018 ± 0.007%/s, r_lit,1_ = –0.309 ± 0.053%/s, r_dark,2_ = –0.043 ± 0.012%/s, r_lit,2_ = –0.229 ± 0.026%/s).

These results demonstrate that the gear-shifting minifilaments can contract actin bundles *in vitro* and that the contraction rates increase reversibly in blue light.

### Optical control of minifilament assembly

Cells use RLC phosphorylation to dynamically regulate minifilament assembly in space and time^1,2^. We therefore considered alternative designs for engineered minifilaments that would enable optical control of minifilament assembly.

Recently, optogenetic strategies have been developed to convert kinesins into larger assemblies to control microtubule organization using light *in vitro*^22,43^. We took inspiration from these strategies to create a myosin light inducible dimer (MyLID) using the improved light-induced dimer (iLID) system^44^. The general design consists of two components: an N-terminal component that contains a myosin motor domain and lever arm fused to iLID LOV2-SsrA, and a C-terminal component that contains SspB-nano fused to an NMII tail domain (Fig. 4b,c and Supp. Fig. 1). The N-terminal motor is monomeric in the dark, and it attaches to the C-terminal tail in the light (Fig. 4d). We predicted that this construct would only be capable of processive motion in the light. This design can incorporate the motor domain from any type of myosin and the tail domain from any myosin II, as well as an engineered or native lever arm. The first version that we tested, MyLID-XI, has an N-terminal component (MyLID-XI-N) that consists of a *Nicotiana* myosin XI motor domain and an alpha-actinin lever arm, and a C-terminal component (MyLID-XI-C) consisting of the human NMIIB tail domain (Fig. 4b).

**Figure 4.**
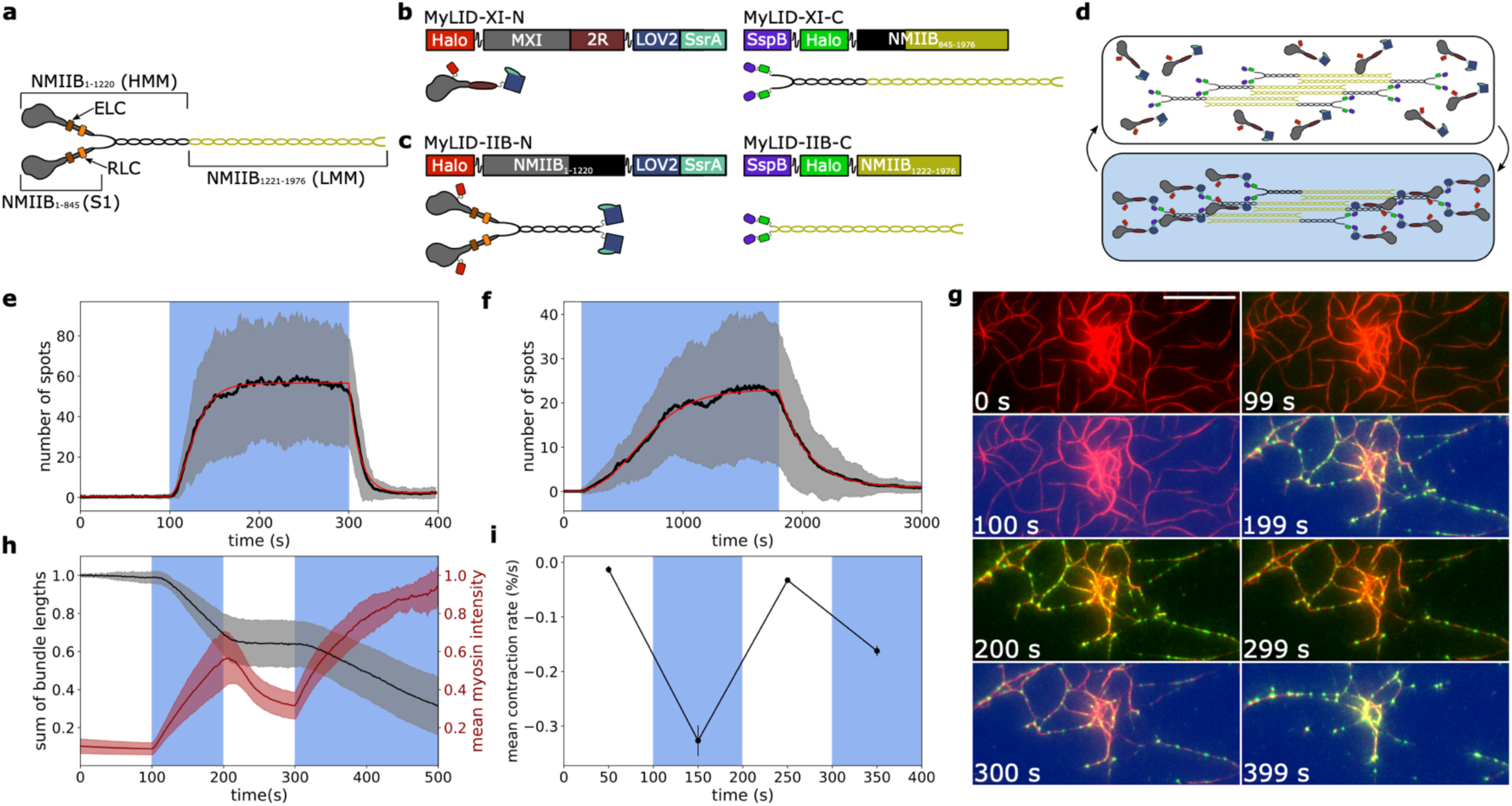
Alternative strategy for optical control of minifilaments using iLID. **a**) Cartoon of a full-length wildtype NMIIB dimer with labels showing different regions of the protein. **b)** Block diagrams and cartoons of MyLID-XI-N (left) and MyLID-XI-C (right). **c)** Block diagrams and cartoons of MyLID-IIB-N (left) and MyLID-IIB-C (right). **d**) Cartoon showing MyLID-XI-N and MyLID-XI-C reversibly assembling into minifilaments in the presence of blue light. **e**) Number of moving spots on the surface over time in a processive motility assay with MyLID-XI on biotinylated actin at 10 μM ATP. MyLID-XI-N and MyLID-XI-C were labeled with 2 different fluorophores, and only MyLID-XI-C was imaged. The black curve is the average of 15 movies across 15 different experiments with 2 protein preps over multiple days, and the gray envelopes represent ± one standard deviation. The red line is a fit to the average curve using a piecewise generalized logistic function rise and exponential decay. Note that these functional forms are empirical fits to the data that are only used to estimate half-lives. **f**) Number of moving spots on the surface over time in a processive motility assay with MyLID-IIB on biotinylated actin at 2 mM ATP in the presence of 0.3% methylcellulose. MyLID-IIB-N and MyLID-IIB-C were labeled with 2 different fluorophores, and only MyLID-IIB-N was imaged. The black curve is the average of 13 movies across 13 different experiments for 1 protein prep over multiple days, and the gray envelopes represent ± one standard deviation. The red line is a fit to the average curve using a piecewise generalized logistic function rise and exponential decay. **g)** Snapshots of a contractility assay with MyLID-XI at 10 μM ATP (taken from Supp. Movie 9) with motors (green) overlayed on top of actin bundles (red). Blue backgrounds indicate that the blue light was on in the given frame. The blue light was off from 0-99 seconds and 200-299 seconds, and left on from 100-199 seconds and 300-500 seconds. The scale bar represents 10 μm. **h)** Sum of the lengths of all actin bundles over time (black curve) and the mean intensity of motors on bundles (red curve) in contractility assays with MyLID-XI at 10 μM ATP. Both curves are normalized to the values in the first frame. Both curves are averages of 21 movies across 10 different experiments with 1 protein prep over multiple days. The envelopes represent ± one standard deviation. **i**) Mean contraction rates in the dark and light for MyLID-XI at 10 μM ATP. Contraction rates were calculated as the slopes of piecewise linear fits to the bundle length vs. time traces. Slopes were calculated for 4 time periods: times 0-99 s (dark), 100-199 s (lit), 200-299 s (dark), and 300-400 s (lit). Each point represents the mean slope (averaged over 21 movies) for each time period. Contraction rates are shown as the percentage change in total bundle length per second. Error bars represent standard errors.

We tested MyLID-XI in processive motility assays on biotinylated actin with fluorophores on the tail domains. MyLID-XI forms puncta during blue illumination that move along actin, while in the dark there are very few motile spots (Supp. Movie 7). In the lit state at 10 μM ATP, the mean velocity is 289 nm/s, and the mean run length is 27.53 μm (Supp. Fig. 7a, Supp. Table 1). We used single particle tracking to calculate the number of motile minifilaments on the surface over time (Fig. 4e). Piecewise rise and decay curve fits to the average number of moving spots on the surface over time give an activation half-life of 28 ± 2 s (mean ± s.e.m) and a deactivation half-life of 10 ± 1 s.

We also sought to create a version of MyLID that mimics the properties of native NMII by using the motor domain and lever arm from wildtype NMIIB. However, previous results suggest that the precise spatial relationship between the 2 motor domains in a myosin II dimer is important for optimal function^45,46^, so we avoided making insertions directly C-terminal to the native lever arm. As an alternative, we inserted the LOV2-SsrA and SspB domains into the NMIIB tail near the heavy meromyosin-light meromyosin junction. We predicted that insertions at this site would not prevent assembly of the tails into minifilaments, as this region was used previously as an insertion site for a force-sensitive FRET module in NMIIB^47^. The resulting construct, MyLID-IIB, has an N-terminal component (MyLID-IIB-N) consisting of native NMIIB up to residue A1220 fused to iLID LOV2-SsrA, and a C-terminal component (MyLID-IIB-C) consisting of SspB-nano fused to the NMIIB tail starting at residue L1222 (Fig. 4c).

We tested MyLID-IIB in assays on biotinylated actin in the presence of 0.3% methylcellulose with fluorophores on the motor domains. MyLID-IIB displays light-dependent recruitment of motile puncta to the surface (Fig. 4d and Supp. Movie 8) with activation and deactivation half-lives of 439 ± 44 s and 255 ± 83 s, respectively. In the lit state at 2 mM ATP, the myosin oligomers walk processively with a mean velocity of 5 nm/s and a mean run length of 2.27 μm (Supp. Fig. 7b, Supp. Table 1). Puncta do not form if only MyLID-IIB-N is included in the assay without MyLID-IIB-C or if MyLID-IIB-N is not phosphorylated (Supp. Fig. 8b,c). The latter observation is consistent with the finding that heavy meromyosin dimers of smooth muscle myosin and NMIIB maintain phosphorylation-dependent activity^36,48,49^. To compare MyLID-IIB to native NMIIB, we also performed assays with wildtype phosphorylated NMIIB minifilaments under the same assay conditions (Supp. Fig. 9). Wildtype NMIIB has a similar mean velocity but a much greater mean run length in comparison to MyLID-IIB (Supp. Table 1), which may be due to a smaller average number of motor domains per myosin oligomer for MyLID-IIB.

To determine if MyLID constructs are capable of contractility, we tested MyLID-XI in contractility assays (Supp. Movie 9). MyLID-XI can contract actin bundles and has a reversible increase in contraction rate in response to blue light (Fig. 4g-i). The average rate of change of the total bundle length increases in magnitude from r_dark,1_ = –0.013 ± 0.007%/s to r_lit,1_ = –0.327 ± 0.028%/s for the first cycle of dark and blue illumination, and from r_dark,2_ = –0.033 ± 0.004%/s to r_lit,2_ = –0.162 ± 0.009%/s for the second cycle (Fig. 4i). The intensity of minifilaments on the bundles also increases reversibly in the lit state (Fig. 4g and Fig. 4h, red curve). In summary, we have shown that MyLID constructs walk processively on actin only in blue light, and they provide an alternative strategy for optical control of contractility *in vitro*.

### Optical control of contractility in live cells

We next applied our engineered minifilaments to control contractile behaviors in live *Drosophila* S2 cells. These cells display a dramatic contractile phenotype in response to activation of their native NMII via the Rho pathway. When plated on concanavalin A (conA)-coated glass, S2 cells spread to form a flattened morphology with a wide lamellar region^50^. After Rho pathway activation, the actomyosin in the lamella contracts, and cells transition to a globular morphology^51^. We predicted that activation of our engineered minifilaments using blue light may stimulate a similar contractile response in these cells.

We expressed MyLOV-XI-IIB labeled with mRuby in S2 cells and used either CellTracker Deep Red or SiR Actin dyes to label the cell morphology. Cells were also treated with the ROCK inhibitor Y27632^52^ to reduce the activity of native NMII. Cells were plated on conA-coated coverslips and imaged in TIRF. Cell morphology and myosin localization varied between transfected cells. MyLOV-XI-IIB was either distributed throughout the lamella or localized to thin cellular protrusions. Cells with protrusions usually did not have the wide, radially uniform lamellae that are typical of cells plated on conA. Similar protrusions were sometimes observed in untransfected cells labeled with SiR Actin. However, protrusions also occurred in CellTracker-labeled cells, indicating that they were not solely due to SiR Actin labeling.

Upon exposure to blue light, cells with well-spread lamellae and high levels of myosin expression in the lamella occasionally displayed radially uniform lamellar contraction around the cell that led to cell shrinkage (Supp. Movies 10 and 11), similar to the contractile phenotype that was described previously^51^, but this behavior was rare. A more common behavior was retraction of protrusions toward the cell body in response to blue light (Supp. Movies 12 and 13), which occurred in all observed cells that displayed both high myosin expression levels and localization of myosin to protrusions. Untransfected cells did not display any blue light-dependent behavior (for example, see the untransfected cells in Supp. Movie 11). We used optical flow of SiR Actin and mRuby-labeled myosin to measure the velocities of protrusions before, during, and after blue illumination (Fig. 5a,b, Supp. Fig. 10). Velocities calculated via optical flow were consistent with velocities estimated from kymographs of retracting protrusions. The average velocity of actin in protrusions increases from 1.4 ± 0.1 nm/s (mean ± s.e.m) before blue illumination to 2.7 ± 0.3 nm/s during illumination and then decreases to 0.8 ± 0.1 nm/s after illumination (with positive velocities indicating motion toward the cell body). The average velocity of myosin in protrusions follows a similar trend (0.3 ± 0.1 nm/s, 2.6 ± 0.4 nm/s, and 0.4 ± 0.1 nm/s before, during, and after blue illumination, respectively). This supports our observation that the protrusion retraction rate increases in the light, and this effect is reversible.

**Figure 5.**
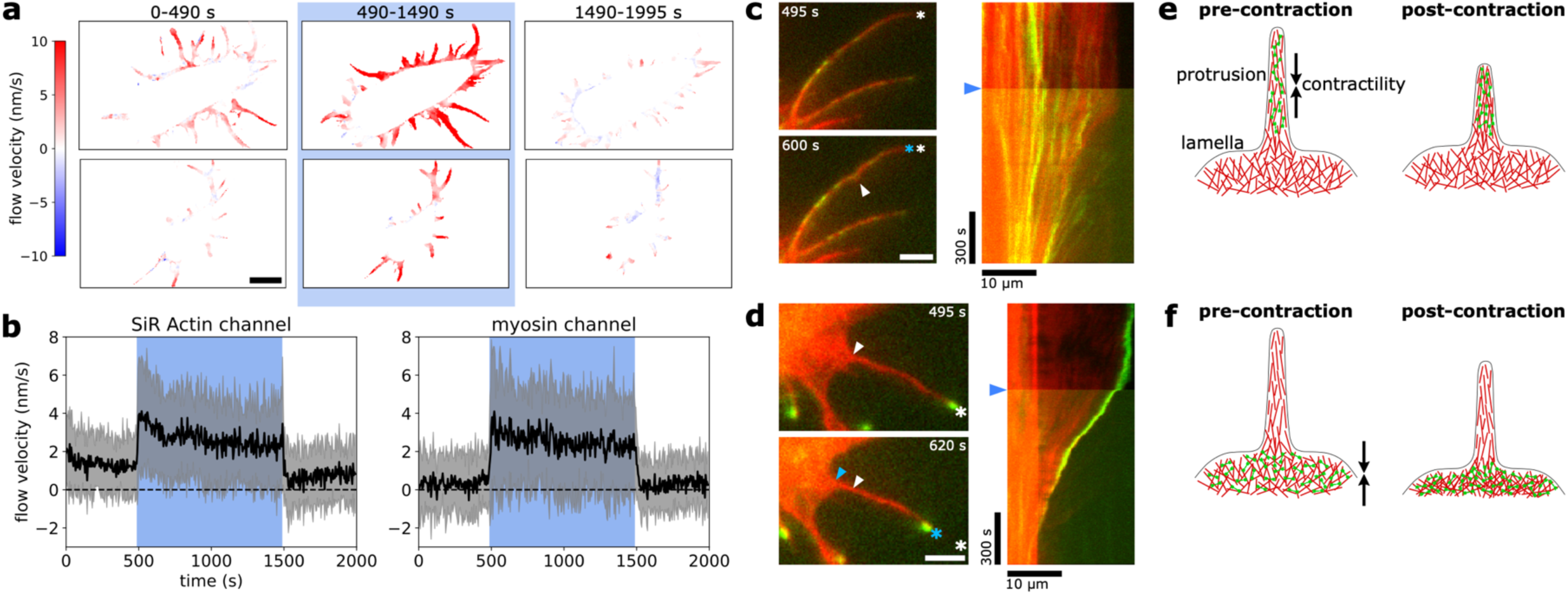
Optical control of protrusion retraction in live S2 cells. **a**) Example SiR Actin mean flow velocity maps for 2 cells. The top cell is shown in Supp. Movie 13. Red/positive indicates flows directed toward the cell body and blue/negative indicates flows directed away from the cell body. Flows were averaged across all frames in the indicated time periods. The blue light was on from 490-1490 s. The scale bar represents 10 μm. **b**) Flow velocity in the protrusions over time for myosin (left) and SiR Actin (right). The black trace is the average across 24 cells imaged over 7 days and the gray envelopes represent ± one standard deviation. **c**) Example of a protrusion undergoing buckling and shortening during blue illumination. The snapshots on the left (taken from Supp. Movie 14) show a protrusion before contraction (top) and during contraction (bottom). Asterisks indicate the position of the protrusion tip (white=position at 495 s, cyan=position at 600 s), showing shortening of the protrusion between the 2 timepoints. The arrowhead indicates a buckle within the protrusion. The scale bar represents 5 μm. A kymograph of the protrusion from 0 to 1490 s is shown on the right with blue arrowhead indicating when the blue light was turned on. **d**) Example of a protrusion being pulled inward as the lamella contracts during blue illumination. The snapshots on the left (taken from Supp. Movie 15) show a protrusion before contraction (top) and during contraction (bottom). Asterisks indicate the position of the protrusion tip, and arrowheads indicate the position of the protrusion base where it connects to the cell body (white=positions at 495 s, blue=positions at 620 s). Simultaneous translation of the protrusion tip and base indicate that it is retracting as an approximately rigid body. The scale bar represents 5 μm. A kymograph of the protrusion from 0 to 1490 s is shown on the right with blue arrowhead indicating when the blue light was turned on. **e)** Cartoon of possible mechanism of retraction of the protrusion shown in **c)**. Myosin (green) localized to the protrusion contracts the actin structures (red) within the protrusion, leading to protrusion shortening. **f)** Cartoon of possible mechanism of retraction of the protrusion shown in **d)**. Myosin localized in the lamella near the base of the protrusion contracts the lamellar actin network. The actin within the protrusion is coupled to the lamellar actin via myosin or other crosslinkers, so the lamellar actin network pulls the protrusion inward toward the cell body.

We observed examples of protrusions buckling and shortening (Fig. 5c, Supp. Movie 14), as well as examples of protrusions moving as approximately rigid bodies while the material near the base of the protrusions contracted (Fig. 5d, Supp. Movie 15). This indicates that there may be two different mechanisms by which the protrusions retract toward the cell body: 1) contraction within the protrusions themselves causes the protrusions to shorten (Fig. 5e), and 2) contraction within the lamellar actin network pulls the protrusions inward as approximately rigid bodies, with myosin or other proteins acting as crosslinkers to couple the lamellar actomyosin to the actin within the protrusion (Fig. 5f). The first mechanism is reminiscent of stress fiber contraction in adherent cells^53^. The second is similar to the proposed mechanism for the retraction of filopodia, in which the filopodium is coupled to the lamella and is dragged inward as NMII contracts the lamellar actin network^54,55^. The lit state retraction velocity is of a similar magnitude to the contraction velocity of stress fibers in fibroblasts during RhoA activation^17^ and the retraction velocity of filopodia in HeLa cells^56^ and COS7 cells^57^. Overall, these results indicate that MyLOV-XI-IIB can contract cellular protrusions and that the contraction rate can be controlled with blue light. However, determining the structure of the actomyosin within the protrusions and the details of the contractility mechanisms will require further investigation.

## Discussion

Here, we have engineered the first examples of filamentous myosins that can be directly controlled by light. We have designed gearshifting minifilaments that change speed and/or direction in response to blue light, cover a wide range of lit state velocities, and walk processively with average run lengths up to several microns. We have also designed minifilaments that can only walk processively on actin in the presence of blue light. Finally, we have demonstrated that the engineered minifilaments can be used for optical control of contractile behaviors both *in vitro* and in cells.

Recent work used engineered myosin multimers to explore the effects of spatially-confined myosin activity^29^ and myosin multimerization state^58^ on the behaviors of extensile actin liquid crystals. Similarly, the engineered minifilaments described here could be used to probe the behaviors of contractile active matter systems. The motor domains and the tail domains of the engineered minifilaments can be varied independently, which enables orthogonal control of biochemical properties and the size of the minifilament. We have also shown here that motor domains from non-filamentous myosins can be used to create minifilaments with novel properties that are not found in native myosin II filaments, such as minus-end-directed motion. This may be useful for investigating how specific microscopic properties of myosin filaments influence macroscopic contractile behaviors, expanding upon previous experimental^7^ and computational^59^ work that examined how contractile behaviors vary across different myosin II isoforms. Overall, both optical control of minifilament activity and modulation of minifilament properties could introduce new types of perturbations into reconstituted systems to improve our understanding of how non-sarcomeric contractility occurs and how contractile systems self-organize.

To our knowledge, we have demonstrated here the first example of a technique for optical control of actomyosin contractility that is both genetically encodable and has been used to control contractility *in vitro*. In contrast, previous techniques either are not genetically encodable^23,24,26,60^ or have only been used in cells^12–17,19,20^ but not *in vitro*. The engineered minifilaments provide a new strategy to introduce spatially and temporally controlled stresses both in living cells and *in vitro,* and results could be compared across both types of experimental systems. This technique may be used to dissect phenomena such as stress fiber contraction and exertion of traction forces^4,11,17^, recruitment of force-sensitive actin binding proteins^17,41,61^, and cortical flows^62,63^. Precisely controlled force generation may be used to probe the material properties of the cytoskeleton^17,64,65^, and comparisons with quantitative physical models of contractile systems^66,67^ will improve our understanding of how nanoscale interactions between cytoskeletal proteins lead to phenomena at the cell and tissue scale.

## Methods

### Molecular constructs and protein purification

Myosin constructs were assembled from DNA fragments using Golden Gate Assembly and cloned into the expression plasmid pBiEx-1 (Novagen). The hybrid minifilaments contained motor domains from *Nicotiana tabacum* myosin XI (codons 1–738), *Sus scrofa* myosin VI (codons 1-816), and *Mus musculus* myosin Va (codons 1-769). The engineered lever arms were assembled from *Dictyostelium discoideum* α-actinin (codons 266-388 for 1R and 267–495 for 2R), *Avena sativa* phot1 (codons 404–543), and *Gallus gallus* α-spectrin (codons 1663–2090). Tail domains were taken from *Homo sapien* nonmuscle myosin IIB (codons 844-1976) and *Drosophila melanogaster* nonmuscle myosin II (codons 849-1964). For the MyLID constructs, the LOV2-SsrA and SspB-nano modules were identical to those used in ref. ^44^ and were fused to other fragments via flexible GSG linkers. The MyLID heterodimers consisted of an N-terminal component with LOV2-SsrA at the C-terminus and a C-terminal component with SspB-nano at the N-terminus. The MyLID-XI head contained the motor domain from *Nicotiana tabacum* myosin XI (codons 1–738) and a lever arm made from *Dictyostelium discoideum* α-actinin (codons 266-502), and the tail contained the *Homo sapien* nonmuscle myosin IIB tail domain (codons 846-1976). MyLID-IIB was assembled entirely from *Homo sapien* nonmuscle myosin IIB (codons 1-1220 for the N-terminal component of the heterodimer and codons 1222-1976 for the C-terminal component). All constructs also contained an N-terminal FLAG tag (DYKDDDDK) and an N-terminal Halotag (Promega) connected to the rest of the protein by a GSG linker. For more details of the construct sequences, see Supp. Fig. 1. The sequences for the full-length wildtype human NMIIB minifilament, chicken essential light chain, chicken regulatory light chain, and myosin light chain kinase (gifts from the Sellers Lab) were all identical to those used in ref. ^32^. M6DI_816_2R∼TET (used for determining the polarity of actin filaments) was identical to the construct M6DI_816_2R∼TET described in ref. ^37^.

Plasmids were directly transfected into Sf9 cells using Escort IV (Sigma) as previously described^68^, or using Fugene HD (Promega). For constructs with native NMIIB lever arms (wildtype NMIIB and MyLID-IIB-N), the myosin plasmid was cotransfected with the essential light chain and regulatory light chain plasmids at an approximately 1:1 molar ratio of myosin plasmid to each light chain plasmid. Bands of the correct size for both light chains were observed on denaturing gels, indicating that the light chains co-purified with the heavy chains.

Proteins were purified from Sf9 cells using anti-FLAG resin as previously described^28^. For constructs with full-length NMII tails, the lysis buffer contained 0.5 M NaCl and the wash buffer contained 0.5 M KCl to prevent the proteins from forming filaments. The proteins were labeled with either tetramethylrhodamine (TMR) or Alexa 660 Halotag ligands (Promega). Proteins were eluted into glycerol storage buffer (500 mM KCl for constructs with full-length tails or 25 mM KCl for all other constructs, 2 mM MgCl_2_, 5 mM EGTA, 25 mM imidazole pH 7.5, 0.1 mM ATP, 1 μg/ml aprotinin, 1 μg/ml leupeptin, 2 mM DTT, and 55% glycerol by volume), aliquoted and flash-frozen in liquid nitrogen within one week after elution, and stored at –80°C. Protein concentrations were measured using SDS-PAGE gel electrophoresis with Sypro Ruby staining. Concentrations were calculated via gel densitometry in Fiji using BSA concentration standards.

### In vitro phosphorylation of native NMIIB constructs

Wildtype NMIIB minifilaments and MyLID-IIB-N, which both contain the native NMIIB lever arm and light chains, were phosphorylated on the regulatory light chain prior to experiments. Phosphorylation buffer consisted of glycerol storage buffer plus 1 μM calmodulin, 500 μM ATP, 5 mM CaCl_2_, 50-100 nM myosin light chain kinase, and 4 mM MgCl_2_. Motors were phosphorylated at room temperature for 30-60 minutes and then stored on ice.

### TIRF Microscopy

All microscopy experiments were performed on a custom-built total internal reflection fluorescence (TIRF) setup with a 1.49 NA, 100x objective (Nikon). Fluorescence was recorded on an electron-multiplying charge-coupled device (EMCCD) camera (Andor iXon DU-897E-CS0-#BV) at maximum gain. Imaging was performed using 3 laser wavelengths: 488 nm (Spectra Physics), 532 nm (Coherent Sapphire), and 633 nm (Blue Sky Research). A light-emitting diode (LED) light source (Thorlabs, M470L3, center wavelength of 470 nm, irradiance of ∼50 mW/cm^2^ at the back focal plane) provided continuous-wave blue light stimulation via the TIRF illuminator arm (Nikon T-FL-TIRF2). The LED was turned on and off manually during imaging. All imaging was performed at room temperature (22-25°C).

### In vitro processive single-molecule motility assays on biotinylated actin filaments

Processive motility assays with full-length minifilaments on biotinylated actin were performed as described in ref. ^32^ with some modifications. Assay buffer consisted of 25 mM Tris (pH 7.5), 25 mM KCl, 1 mM EGTA, 2 mM MgCl_2_, and 10 mM DTT. Blocking buffer consisted of 25 mM Tris (pH 7.5), 25 mM KCl, 1 mM EGTA, 2 mM MgCl_2_, 10 mM DTT and 2 mg/ml BSA (Sigma-Aldrich). Imaging buffer consisted of blocking buffer supplemented with ATP, an oxygen-scavenging system (0.2 mg/ml glucose oxidase, 0.36 mg/ml catalase, and 0.4% wt/vol glucose), and an ATP regeneration system (0.95 μg/ml creatine phosphokinase and 1 mM phosphocreatine). ATP concentrations were 500 μM for MyLOV-XI-IIB, MyLOV-XI_L2+4_-IIB, and MyLOV-V-IIB, 2 mM ATP for MyLOV-VI-IIB, MyLOV-VI_L310G_-IIB, MyLOV-VI-*zip*, MyLID-IIB, and wildtype NMIIB, and 10 μM for MyLID-XI. For constructs with myosin VI heads, 1 μM calmodulin was also added to the imaging buffer. For MyLID-IIB and wildtype NMIIB minifilaments, 0.3% methylcellulose was included in the imaging buffer. Minifilaments were formed by rapidly diluting the motors from high-salt storage buffer into low-salt imaging buffer right before adding the imaging buffer to the channel. Motor concentrations were adjusted to achieve optimal motor density for single particle tracking. The final motor concentration varied from ∼300 pM to ∼5 nM depending on the construct. To determine the directionality of the actin filaments, the unidirectional motor M6DI_816_2R∼TET was included in the imaging buffer. For the constructs MyLOV-XI-IIB, MyLOV-XI_L2+4_-IIB, and MyLOV-V-IIB, we observed that close to 100% of the runs are plus-end-directed in the lit state (Supp. Table 2). Most data for these 3 constructs was therefore collected without using M6DI_816_2R∼TET to determine the actin polarity, and the polarity of each actin filament was instead determined by the direction that the minifilaments moved in the lit state. Biotinylated actin was prepared at a 5% concentration of biotinylated monomers by mixing biotinylated G-actin (Cytoskeleton Inc.) with unlabeled G-actin (gift from S. Sutton). Actin filaments were labeled with Alexa 488 phalloidin (Invitrogen).

Assays were performed in flow channels assembled from a nitrocellulose-coated coverslip attached to a slide using double-sided tape, as described in ref. ^28^. Channels were prepared for imaging using the following steps (with 2-5 minutes of incubation between each step): 1) 1 mg/ml biotinylated BSA (ThermoFisher), 2) 1 mg/ml neutravidin (Invitrogen), 3) blocking buffer, 4) biotinylated actin diluted in assay buffer, and 5) imaging buffer. The channels were then sealed with vacuum grease.

Each movie consisted of at least one period of dark and one period of blue LED illumination lasting at least 100 s each. The minifilaments (labeled with TMR Halotag ligand) were imaged first with the 532 nm laser (50-100 μW at the back aperture), with an exposure time ranging between 0.3 s and 3 s depending on the construct. For the MyLID constructs, the N-terminal and C-terminal components were labeled with different fluorophores (either Alexa 660 or TMR) to distinguish between the 2 components, and only the TMR fluorophores were used for imaging. For MyLID-XI, only MyLID-XI-C was imaged, and for MyLID-IIB, only MyLID-IIB-N was imaged. For experiments in which polarity scoring of actin was required, the myosin VI tetramers (labeled with Alexa 660 Halotag ligand) were imaged with the 633 nm laser for 5-10 minutes after imaging the minifilaments. The actin was imaged with the 488 nm laser for a few frames after imaging the motors.

Motor velocities and run lengths were measured by analyzing kymographs of motor trajectories along single actin filaments. Actin filaments were traced in ImageJ and kymographs were produced via the KymographBuilder plugin. The start and end of each individual motor run were selected manually, and the run was then traced using the ImageJ Single Neurite Tracker (SNT) plugin^69^. ROIs of the traced trajectories were saved as text files and then analyzed in Python to calculate velocities and run lengths. Runs that were observed by eye to have 0 velocity and to remain attached to actin during the entire movie were assumed to be irreversibly stuck to actin and were not measured. To further filter out stuck motors, measured runs were rejected during analysis if the total displacement was <= 320 nm (2 pixels) and the run was >= 200 seconds long. Each run was identified as either censored (ran off the actin filament, was cut off at the end of the movie, or was cut off during a transition to a different light state) or uncensored (dissociated from the actin filament). Run censoring was determined automatically for MyLOV-XI-IIB, MyLID-IIB, and wildtype NMIIB by calculating the distance from the last position in the run to the end of the distance axis of the kymograph. For all other constructs, censoring was determined manually because runs frequently merged or crossed over each other. The velocity of each run was calculated as total displacement divided by total time. The time-weighted average of the velocities of all runs was calculated for a single experiment, and the final mean velocity was calculated by averaging across all experiments. The standard error of the mean velocity was calculated across experiments. The mean run length was calculated as the sum of all run lengths across all experiments divided by the total number of uncensored runs, which is the maximum likelihood estimate of the mean for a right-censored exponential distribution. The standard error of the mean run length was calculated using bootstrapping with 1000 samples. The mean run length was not calculated if >90% of all runs were censored.

For MyLID-XI and MyLID-IIB, the number of motors on the surface over time was calculated by performing single particle tracking using the Trackmate plugin in ImageJ^70^. Analysis of tracks was performed in Python. To filter out stuck and diffusing particles, tracks with fewer than 3 timepoints or with displacement < 640 nm (4 pixels) were rejected during analysis. In addition, a linear fit was performed for log(mean squared displacement) vs. log(time) for each track and the slope of the line was used to distinguish between stuck/diffusing particles and particles that were moving along actin. The minimum threshold for the slope was 1.0 for MyLID-IIB and 1.2 for MyLID-XI. Traces of number of moving motors on the surface vs. time were calculated for each movie and then averaged across multiple movies for each construct (15 movies for MyLID-XI, 13 movies for MyLID-IIB). To estimate rise and decay half-lives, piecewise nonlinear fitting of the traces was performed using the curve_fit function from scipy.optimize. Rise portions of the curve were fit to a generalized logistic function, and decay portions were fit to an exponential function. Half-lives were calculated for each individual experiment and then averaged across all experiments to compute mean half-lives.

### In vitro processive single-molecule motility assays on fascin-actin bundles

Motility assays on fascin-actin bundles were adapted from a protocol described in ref. ^40^. Assay buffer, blocking buffer, and imaging buffer were the same as those used in motility assays on biotinylated actin, except that the KCl concentration in the imaging buffer was increased from 25 mM KCl to 75 mM KCl. We observed that minifilaments on bundles frequently collide and merge into large clusters in low salt, and increasing the salt concentration prevents this clustering behavior. Actin filaments were grown from gelsolin-capped seeds (see ref. ^71^ for detailed protocol) and labeled with a either mixture of dark phalloidin and Alexa 488 phalloidin or only dark phalloidin. Fascin-actin bundles were formed by incubating actin filaments at a concentration of 1.7 mg/ml actin monomers with 20 μg/ml recombinant human fascin (Abcam ab99240). The fascin-actin mixture was incubated at room temperature for 5-10 minutes before adding to the channel.

Assays were performed in flow channels identical to those used in motility assays on biotinylated actin. Aligned and polarized actin bundles were formed using the following steps (with 2-5 minutes of incubation between each step): 1) anti-gelsolin (Sigma G4896) diluted 1 in 7 in assay buffer, 2) blocking buffer, 3) the mixture of fascin and actin described above, 4) blocking buffer supplemented with 8 μg/ml fascin, and 5) imaging buffer with 8 μg/ml fascin, 2 mM ATP, and 1 μM calmodulin. The channels were then sealed with vacuum grease. Flow steps 3-5 were performed as quickly as possible using capillary action. The anti-gelsolin on the surface binds to the gelsolin within the bundles, and the rapid flow causes the free ends of the bundles to align along the flow direction and flatten against the surface^40^. As a result, most bundles have their plus-ends facing the entrance of the channel (where buffers were flowed in) and their minus-ends facing the exit of the channel. The bundles likely remain flattened against the surface by a combination of gelsolin/anti-gelsolin interactions and fascin cross-linking between different bundles. Fascin must be used in all flow steps after adding the bundles to the channel, or else the bundles will detach from the surface.

Actin bundles were briefly imaged first using 488 nm laser excitation when Alexa 488 phalloidin was used, and then the motors (labeled with TMR) were imaged using 532 nm laser excitation. Motors were imaged at 2 s per frame with a 200 s period of dark, followed by a 200 s period of blue LED illumination.

Motor velocities were calculated using kymograph analysis, as described above for assays on biotinylated actin, with some modifications (see Supp. Fig. 5). It was difficult to define individual bundles for the purpose of generating kymographs, as the assay requires the bundles to be packed tightly together, and bundles frequently merge together and branch apart. To generate the kymographs, continuous paths of motors along bundles were traced using a maximum intensity projection of the motors. Run lengths were not calculated.

### Contractile bundle assays

Assay buffer was the same as that used in the processive motility assays. Blocking buffer and imaging buffer were the same except for the addition of 0.3% w/v methylcellulose in both. Actin bundles were formed by incubating actin filaments (labeled with Alexa 633 phalloidin) at a concentration of 20 μg/ml with 15 μg/ml alpha-actinin (Cytoskeleton Inc.). The alpha-actinin-actin mixture was incubated at room temperature for 5-10 minutes before adding to the chamber.

Assays were performed in an open chamber to avoid fast flows (which could disturb the actin network after it has formed) and to enable addition of motors to the chamber during imaging. Cylindrical chambers were formed from 0.2 ml PCR tubes by removing the cap and the bottom 2/3 portion of the tube. The remaining 1/3 top portion of the tube was attached to a nitrocellulose-coated coverslip using nail polish. A network of actin bundles was formed using the following steps (with 2-5 minutes of incubation between each step): 1) 20 μl of 10 mg/ml BSA (Sigma), spread with a pipette tip to ensure complete coverage of the surface of the chamber, 2) 10 μl of the mixture of actin and alpha-actinin described above, plus 5 μl of blocking buffer, and 3) 25 μl of imaging buffer without motors and supplemented with 3 μg/ml alpha-actinin. Before steps 2 and 3, the solution from the previous step was removed by pipetting as slowly as possible, and a few μl of liquid was left in the chamber.

The actin bundles were observed first using 633 nm laser excitation before the motors were added and a region with optimal bundle architecture was located for imaging. The motors (labeled with TMR) were then added to the chamber and observed using 532 nm laser excitation. The final motor concentrations were 100-300 pM for MyLOV-VI-IIB and MyLOV-XI-IIB and 10 nM for MyLID-XI. For MyLOV-VI-IIB and MyLOV-XI-IIB, the motors were monitored until there was an approximately constant density of motors on the bundles (which usually occurred a few minutes after the motors were added), and then the actin was imaged. For MyLID-XI, the motors and actin were imaged simultaneously using an Optosplit. Movies were acquired at 1 s per frame with the following illumination sequence: 100 s dark, 50 s blue illumination (for MyLOV-VI-IIB and MyLOV-XI-IIB) or 100 s blue illumination (for MyLID-XI), 100 s dark, then blue illumination until contraction was no longer observed (usually 200-300 s).

Contraction rates over time were measured by calculating the sum of the lengths of all actin bundles within the field of view in each frame (see Supp. Fig. 7). The analysis was performed in Python. First, each frame of a movie was thresholded using the local threshold function from scikit-image. Next, the thresholded images were skeletonized using the skeletonize function from scikit-image. To reduce noise, branches less than 5 pixels long were removed from the skeletonized images using the Skeleton Analysis (skan) package. Finally, the total bundle length for each image was measured as the total number of pixels in the skeleton. Contraction rates for each period of dark and each period of blue illumination in one movie were measured as the slope of the total bundle length vs. time trace. Slopes were calculated by performing linear fits for each period of dark and illumination. For the second period of illumination, only the first 50 s (for MyLOV-VI-IIB and MyLOV-XI-IIB) or 100 s (for MyLID-XI) were used to calculate the contraction rate. For each construct, the contraction rates were calculated for each movie and then averaged across all movies to compute mean contraction rates.

### S2 cell culture and molecular constructs for live cell experiments

*Drosophila* S2 cells were purchased from Gibco. Cells were grown in tissue culture dishes at room temperature in Schneider’s *Drosophila* Medium (Gibco) with 10% heat-inactivated fetal bovine serum (Sigma). The coding sequence of MyLOV-XI-IIB without the N-terminal Halotag was transferred from the pBiEx-1 vector to the pMT *Drosophila* expression vector and fused to an N-terminal mRuby tag using Gibson Assembly. The pMT backbone was derived from the plasmid pMT-EGFP-Actin5c (gift from Ron Vale, Addgene plasmid #15312). Transfections were performed using a Lonza 4D nucleofector with SF Cell Line solution and the DG-135 nucleofection protocol following the manufacturer’s instructions. Cells were induced with 500 μM or 1 mM copper sulfate between 6 and 24 hours after transfection. Imaging was performed between 2 and 5 days after induction.

### Live cell TIRF imaging

Cells were stained with either CellTracker Deep Red (Invitrogen) or SiR Actin (Cytoskeleton Inc.) prior to imaging. For staining with CellTracker Deep Red, cells were incubated in a solution of 300 nM of CellTracker for one hour. The staining solution was replaced with Schneider’s Medium without stain for imaging. For staining with SiR Actin, cells were incubated in a solution of 1 μM SiR Actin + 10 μM Verapamil for one hour. Cells were then washed with media and the staining solution was replaced with a solution of Schneider’s Medium + 100 nm SiR Actin + 10 μM Verapamil for imaging. In addition, cells were incubated with 20 μM Y27632 for at least one hour before imaging, and 20 μM Y27632 was included in the media during imaging.

Concanavalin A (conA) coated coverslips were prepared by coating plasma-cleaned coverslips with a 0.5 mg/ml solution of conA (Sigma) and allowing them to air dry overnight. On the day of imaging, additional conA solution was added to the imaging chambers and incubated for 15-30 minutes. Cells were allowed to spread on the conA-coated coverslip for at least one hour prior to imaging.

Cells were simultaneously imaged using both 633 nm excitation to visualize CellTracker Deep Red or SiR Actin and 532 nm excitation to visualize mRuby-labeled myosin. Movies were acquired at 5 s per frame with the following blue illumination sequence: 490 s of dark, 1000 s of blue illumination, and 505 s of dark. Only cells labeled with SiR Actin were used for analysis, as CellTracker Deep Red usually did not label the protrusions. Only cells that displayed both SiR Actin labeling of protrusions and high levels of myosin expression were included in the analysis. Occasionally cells with protrusions also displayed highly active lamellae that underwent cycles of extension and retraction. These cells were excluded from the analysis because the lamellar motion added noise to the optical flow measurements. The final dataset contained 24 cells imaged across 7 days.

To calculate protrusion velocities, first a mask was created manually from the mean projection of the SiR Actin channel to remove the cell body and most of the background. The protrusions were then identified in each frame in the SiR Actin channel using ridge detection (scikit-image meijering filter) and all non-ridge pixels were excluded to create a final mask of the protrusions. Flow velocities were calculated in both the SiR Actin and myosin channels using Farneback optical flow in opencv. The protrusion mask was used to identify the flow vectors corresponding to the locations of the protrusions in each frame, and only these flow vectors were included in the analysis. To convert flow vectors into flow velocities inward/outward relative to the cell body, the approximate position of the cell centroid was determined by calculating the centroid of the cell body in the mean projection of the SiR Actin channel. Each flow vector was then projected onto the vector pointing from the flow vector coordinate to the cell centroid coordinate. Traces of flow velocity vs. time were calculated for each cell by averaging the projected flow velocities in each frame. Mean flow velocities were calculated by averaging the flow velocities over time for each cell within each light period (0-490 s for pre-illumination, 495-1490 s for during illumination, and 1495-1995 s for post-illumination) and then taking the mean across 24 cells for each light period.

## Acknowledgements

This work was supported by an NSF Graduate Research Fellowship to S.Z., NIH awards R01GM114627 and R01GM143792, and NSF award DMR-2215605. We thank Margaret Gardel, Jordan Beach, Patrick Oakes, Stephan Grill, Enrico Perini, Gregory Alushin, and members of the Bryant lab for useful discussions and exchanges.

