## Supplementary Material for "Engineering filamentous myosins for optical control of contractility"

| construct | state | [ATP] (µM) | mean velocity (nm/s) | modulation depth | mean run length (µm) | fraction censored runs | # of experiments | total # of runs |
|---|---|---|---|---|---|---|---|---|
| MyLOV-XI-IIB | dark | 500 | 50 ± 6 | 0.89 ± 0.01 | 0.73 ± 0.02 | 0.352 | 12 | 1741 |
| | lit | | 497 ± 25 | | 2.21 ± 0.05 | 0.444 | | 2941 |
| MyLOV-$XI_{L2+4}$-IIB | dark | 500 | 17 ± 1 | 0.80 ± 0.01 | N.D. | 0.996 | 10 | 1161 |
| | lit | | 82 ± 3 | | N.D. | 0.998 | | 1247 |
| MyLOV-V-IIB | dark | 500 | 11 ± 1 | 0.49 ± 0.03 | N.D. | 0.963 | 10 | 630 |
| | lit | | 22 ± 1 | | N.D. | 0.964 | | 633 |
| MyLOV-VI-IIB | dark | 2000 | 4 ± 1 | 1.07 ± 0.02 | N.D. | 0.976 | 10 | 1191 |
| | lit | | -63 ± 3 | | N.D. | 0.990 | | 1774 |
| MyLOV-$VI_{L310G}$-IIB | dark (BT) | 2000 | 18 ± 1 | 1.16 ± 0.01 | 7.15 ± 0.47 | 0.799 | 10 | 1162 |
| | lit (BT) | | -112 ± 4 | | 22.29 ± 1.63 | 0.885 | | 1662 |
| | dark (fascin) | | 28 ± 4 | 1.36 ± 0.07 | N.D. | N.D. | 10 | 3182 |
| | lit (fascin) | | -89 ± 6 | | N.D. | N.D. | | 4192 |
| MyLOV-VI-*zip* | dark | 2000 | 12 ± 1 | 1.26 ± 0.03 | N.D. | 0.991 | 10 | 1018 |
| | lit | | -52 ± 5 | | N.D. | 0.995 | | 1316 |
| MyLID-XI | lit | 10 | 289 ± 5 | N.A. | 27.53 ± 2.50 | 0.868 | 4 | 955 |
| MyLID-IIB | lit | 2000 | 5 ± 0.3 | N.A. | 2.27 ± 0.13 | 0.434 | 13 | 466 |
| WT NMIIB | N.A. | 2000 | 8 ± 0.4 | N.A. | 18.19 ± 2.52 | 0.886 | 13 | 519 |

**Supplementary Table 1. Summary of velocity and run length measurements for all constructs in processive motility assays.** For MyLOV-$VI_{L310G}$-IIB, separate measurements are shown for assays on biotinylated actin and assays on fascin-actin bundles, which are indicated with (BT) and (fascin) respectively. Mean run lengths were calculated using the maximum likelihood estimate of the mean for right-censored data from an exponential distribution. N.D. = not determined; mean run lengths were not calculated for constructs with fraction censored runs >0.9. Run lengths were not calculated for assays on fascin-actin bundles. Measurements are shown as mean ± standard error.

| construct | state | # of plus-end-directed runs | # of minus-end-directed runs |
|---|---|---|---|
| MyLOV-XI-IIB | dark | 512 | 14 |
| | lit | 750 | 1 |
| MyLOV-$XI_{L2+4}$-IIB | dark | 243 | 1 |
| | lit | 295 | 1 |
| MyLOV-V-IIB | dark | 139 | 1 |
| | lit | 112 | 0 |

**Supplementary Table 2. Directionality of myosin XI and myosin V constructs in processive motility assays on biotinylated actin.** Actin polarity was determined by imaging MVI-2R-tet after imaging the minifilaments. Runs were counted manually from kymographs. Data was collected from 4 separate experiments with 2 protein preps for each construct. Assays were performed at 500 µM ATP for MyLOV-$XI_{L2+4}$-IIB and MyLOV-V-IIB and at 100 µM ATP for MyLOV-XI-IIB.

Wildtype NMIIB
DYKDDDDKMAEIG...EISGEPTTEDLYFQSDNAARGIQMAQRT...PQSE
NMIIB 1-1976

MyLOV-XI-IIB
DYKDDDDKGGSSEIG...PGLAGSASVN...GNAAQTRS...KIAVLATT...DEAATRSD...AELENKQQ...TLVAGSLLQV...PQSE
MXI 1-738

MyLOV-XI$_{L2+4}$-IIB
DYKDDDDKGGSSEIG...PGLAGSASVN...LPEES**GGKKGG**SKSSK**KK**FSS...GNAAQTRS...KIAVLATT...DEAATRSD...AELENKQQ...TLVAGSLLQV...PQSE
MXI loop 2 insertion

MyLOV-V-IIB
DYKDDDDKGGSSEIG...PGLAGSAASE...LRAAQTRS...KIAVLATT...DEAATRSD...AELENKQQ...TLVAGSLLQV...PQSE
MVa 1-769

MyLOV-VI$_{L310G}$-IIB
DYKDDDDKGGSSEIG...PGLAGSEDGK...LKDP**G**LDDH...RAEAQTRS...KIAVLATT...DEAATRSD...AELENKQQ...TLVAGSLLQV...PQSE
MVI L310G mutation

MyLOV-VI-*zip*
DYKDDDDKGGSSEIG...PGLAGSEDGK...RAEAQTRS...KIAVLATT...DEAATRSD...AELENKQQ...TLVAGSPLLE...EGLH
*zipper*
849-1964

MyLOV-VI-IIB
DYKDDDDKGGSSEIG...PGLAGSEDGK...RAEAQTRS...KIAVLATT...DEAATRSD...AELENKQQ...TLVAGSLLQV...PQSE
MVI 1-816 | α-actinin 266-388 | AsLOV2 404-543 | α-actinin 267-495 | α-spectrin 1663-2090 | NMIIB 844-1976

MyLID-XI-N
DYKDDDDKGGSSEIG...PGLAGSASVN...GNAAQTRS...KIED(GSG)$_3$LATT...ENYF
MXI 1-738 | α-actinin 266-502

MyLID-XI-C
EYKS...LNIEGSDYKDDDDKGGSSEIG...PGLAGSGQVTR...PQSE
NMIIB 846-1976

MyLID-IIB-N
DYKDDDDKMAEIG...EISGEPTTEDLYFQSDNAARGIQMAQRT...RFKAGSGLATT...ENYF
Halotag | linker | NMIIB 1-1220 | LOV2-SsrA

MyLID-IIB-C
EYKS...LNIEGSDYKDDDDKGGSSEIG...PGLA(GSG)$_2$LEKN...PQSE
SspB-nano | linker | FLAG tag | linker | Halotag | linker | NMIIB 1222-1976

**Supplementary Figure 1. Construct sequences.** Junction and sequence information for the constructs produced in this study. Residue insertions and substitutions are indicated in teal. Flexible linkers between domains are indicated in pink. Subscripts indicate the number of repeats within the linker (e.g. (GSG)$_3$ represents 3 GSG repeats).

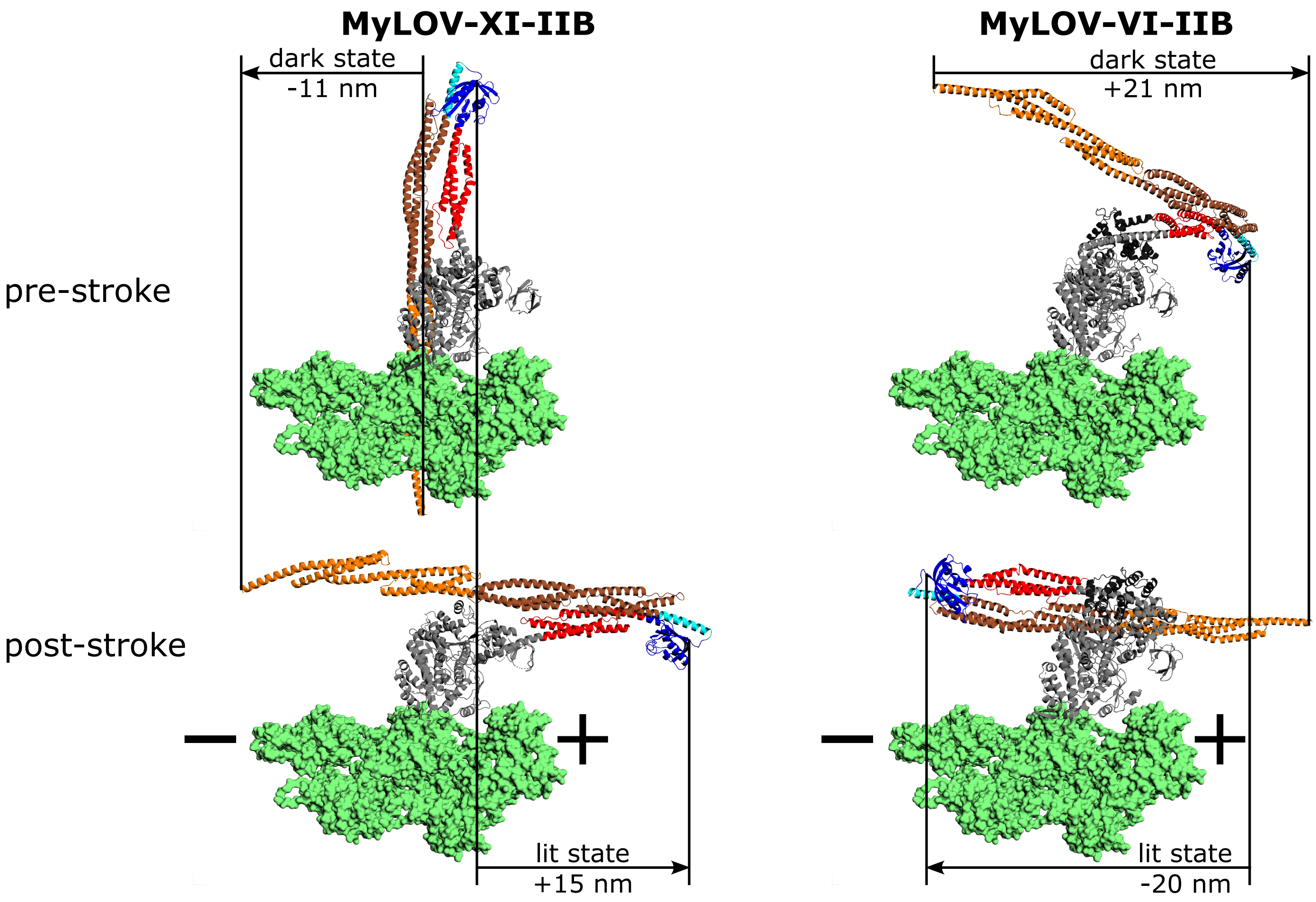

**Supplementary Figure 2. Structural models and predicted stroke vectors for myosin XI and myosin VI constructs.** Lowest-energy structures generated by Rosetta Remodel for the motor domain and lever arm of MyLOV-XI-IIB (left) and MyLOV-VI-IIB (right) in the pre-stroke state (top) and the post-stroke state (bottom). The labels indicate the average stroke lengths in the dark state (top arrows) and lit state (bottom arrows). Stroke lengths were calculated by computing the stroke vector between the tip of the pre-stroke lever arm and the tip of the post-stroke lever arm, and then projecting the stroke vector onto the actin filament. The last residue of the final spectrin repeat was used as the tip of the lever arm in the dark state. The last residue in the LOV2 domain before the hinge that connects to the Jα helix (G516) was used as the tip of the lever arm in the lit state. Average stroke lengths were calculated across the 50 lowest-energy pre-stroke and post-stroke structures. The mean ± standard deviation stroke lengths are: -11 ± 4 nm (dark state) and 15 ± 2 nm (lit state) for MyLOV-XI-IIB, and 21 ± 3 (dark state) and -20 ± 1 (lit state) for MyLOV-VI-IIB.

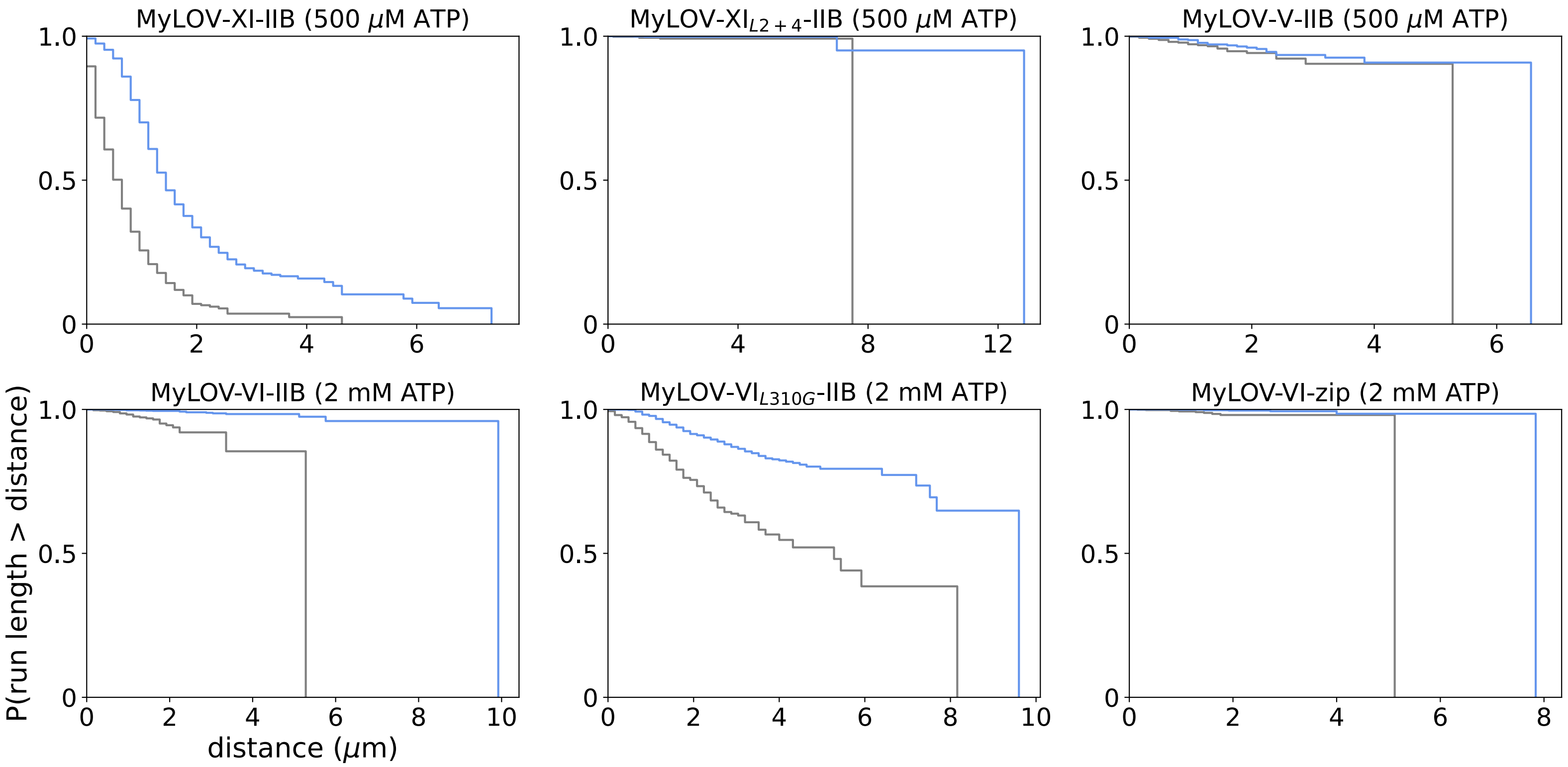

**Supplementary Figure 3. Run length survival curves for gear-shifting minifilaments in processive motility assays on biotinylated actin.** Probability that a run length will be longer than a given distance is plotted for each gear-shifting construct in the dark (gray) and in the presence of blue light (blue). Survival curves were constructed from the same datasets that were used for velocity analysis in Figure 2c and 2d. Run lengths and censoring were determined from kymograph analysis.

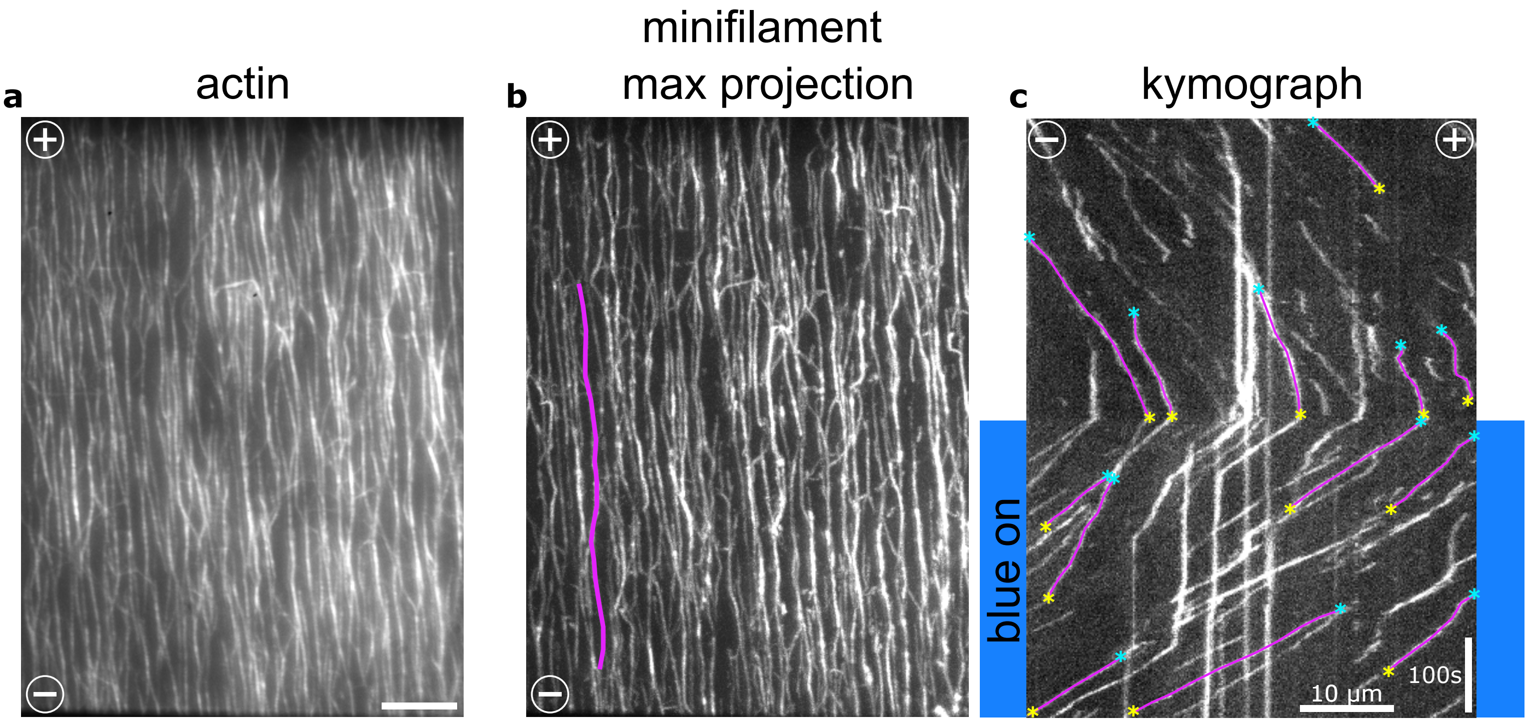

**Supplementary Figure 4. Analysis procedure for processive motility assays on fascin-actin bundles. a)** Fluorescence image of aligned and polarized fascin-actin bundles. Plus-ends are oriented toward the top of the image and minus-ends are oriented toward the bottom. Scale bar represents 10 µm. **b)** Maximum intensity projection of the motor channel in the same field of view as shown in **a)**. An example of a traced bundle is labeled in magenta. **c)** Kymograph generated from the traced bundle shown in **b)**. The minus-end of the bundle is on the left side of the kymograph and the plus-end is on the right. Examples of individual runs that would be analyzed for velocity calculations are labeled in magenta, with the starting points of the runs labeled with cyan asterisks and the ending points labeled with yellow asterisks. The blue background indicates the period of blue illumination beginning at 400 seconds.

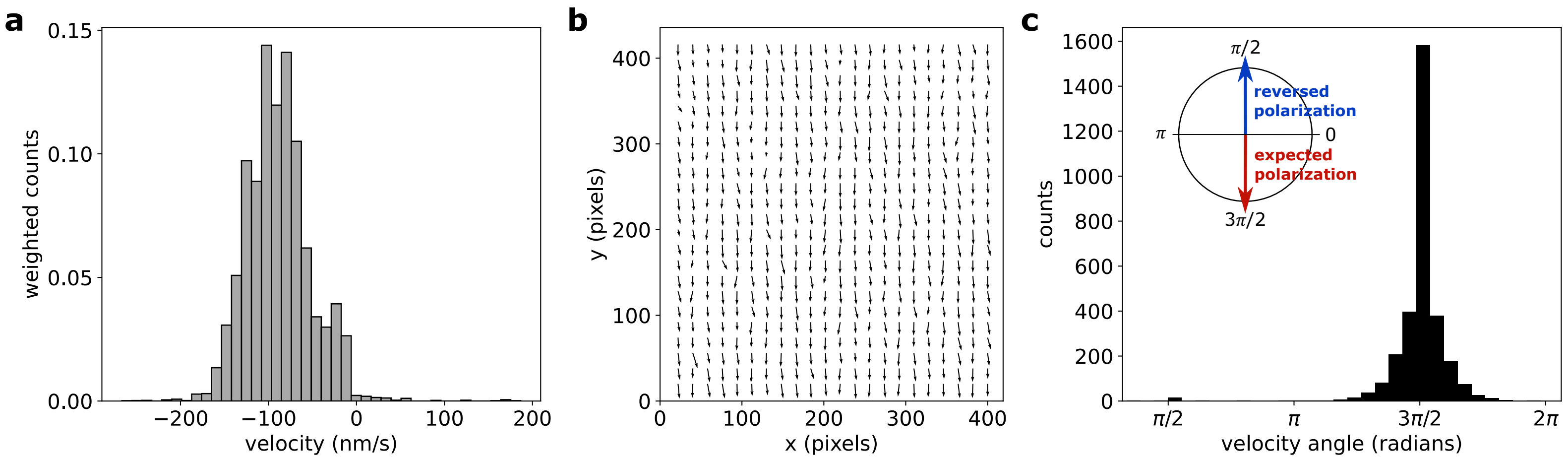

**Supplementary Figure 5. Polarization of aligned fascin-actin bundles. a)** Histogram of run velocities for a unidirectional myosin VI tetramer on aligned fascin-actin bundles. Positive velocities indicate runs that were scored the with incorrect polarity (~1% of all runs), while negative velocities were scored with the correct polarity. The mean velocity is -92±9 nm/s (mean ± s.e.m.). The data was collected from 3 separate experiments. **b)** Average velocity field for the myosin VI tetramers on fascin-actin bundles calculated from PIV analysis. The data was collected from 6 total movies across 3 different experiments and 6 different fields of view. The time-averaged velocity field was calculated for each movie, and the average was taken across all movies to produce the final average velocity field. In this coordinate system, actin bundles are expected to be polarized along the y-direction, with plus-ends pointing in the +y direction and minus-ends pointing in the -y direction. The motor velocity field indicates that most bundles are polarized in this orientation. **c)** Histogram of the angles of time-averaged velocity vectors for all movies calculated from PIV analysis using the same dataset as in **b)**. An angle close to 3π/2 indicates that the actin bundle along which the motor is moving is polarized in the expected orientation, while an angle close to π /2 indicates that the polarity is reversed. The angle distribution indicates that most bundles are polarized in the expected orientation.

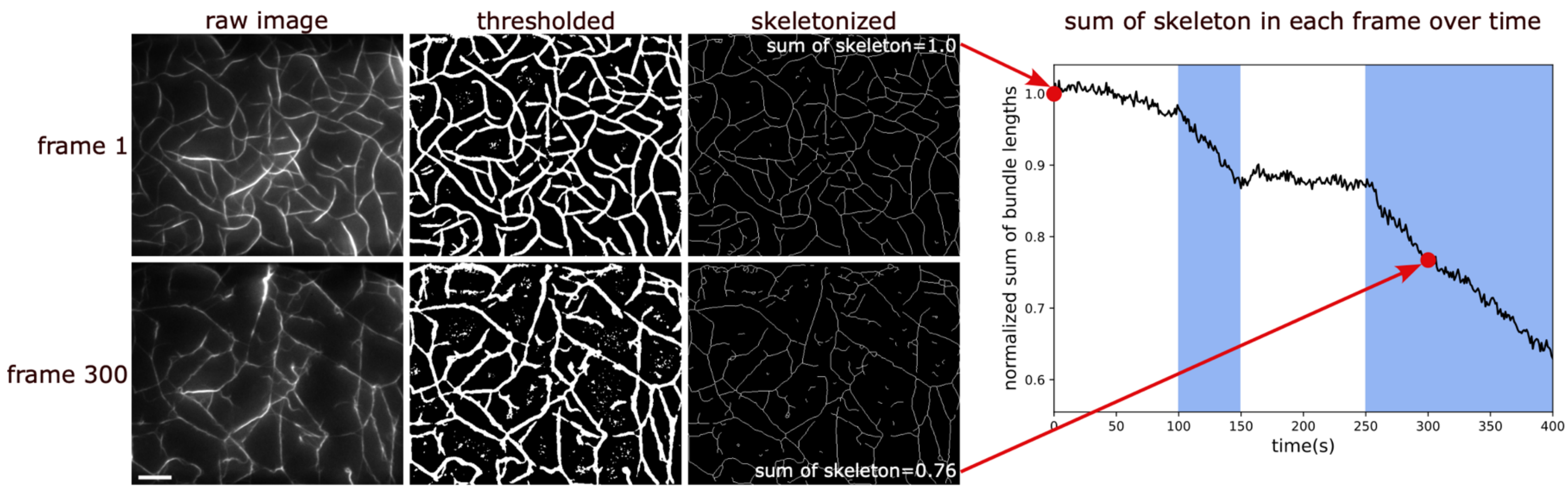

**Supplementary Figure 6. Analysis procedure for *in vitro* contractility assays.** Example analysis for the 2 snapshots from the contractility assay with MyLOV-VI-IIB shown in Figure 3b. The raw fluorescence image of the bundles of actin and α-actinin (left) is thresholded (center) and then skeletonized (right). The total bundle length in each frame is computed as the sum of all pixels in the skeleton and normalized to the sum of the skeleton in frame 1. The normalized sum of the skeleton in each frame then contributes to one timepoint in a plot of the sum of bundle lengths over time, as shown on the right.

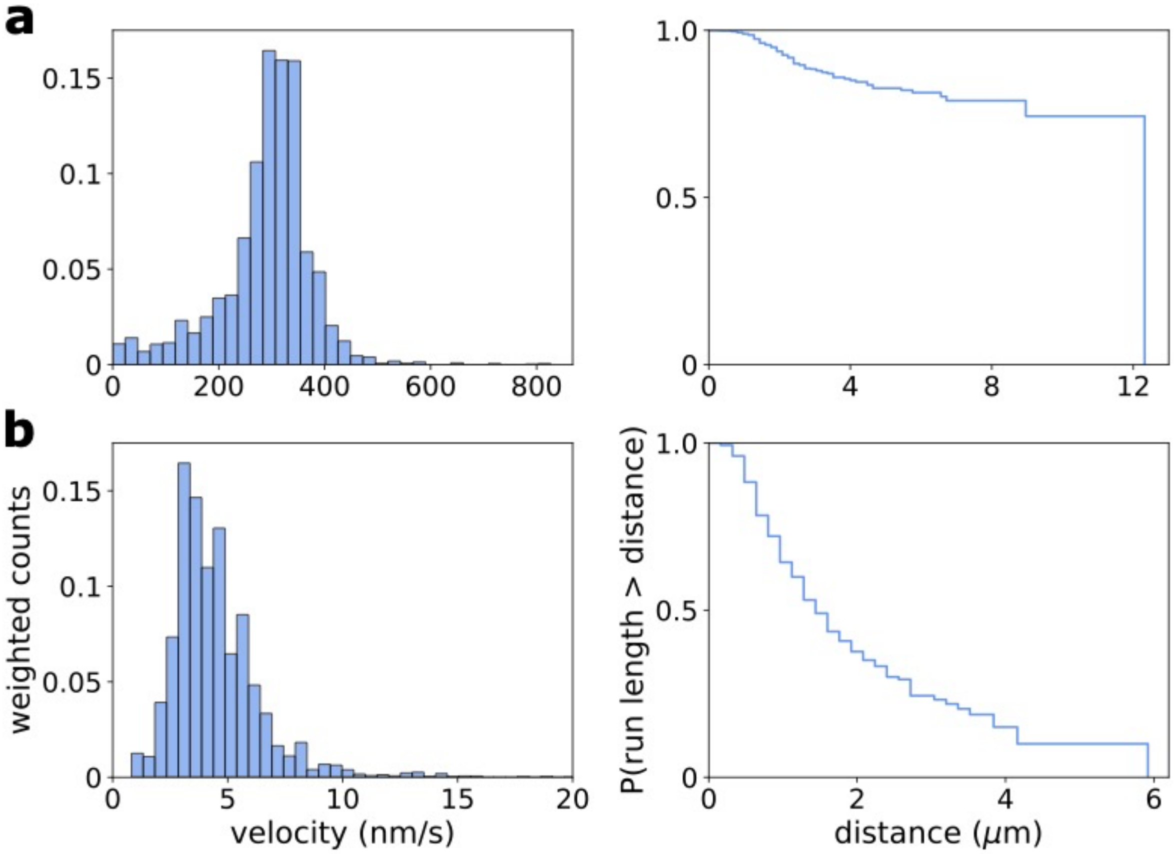

**Supplementary Figure 7. Velocities and run lengths of MyLID constructs in processive motility assays on biotinylated actin. a)** Velocity histogram (left) and run length survival curve (right) for MyLID-XI at 10 μM ATP. Data was collected from 4 separate experiments with 2 protein preps over 2 days. **b)** Velocity histogram (left) and run length survival curve (right) for MyLID-IIB at 2 mM ATP in the presence of 0.3% methylcellulose. Velocity values above 20 nm/s (~0.2% of all runs) were cut off in the histogram for data visualization. Data was collected from 13 experiments with 1 protein prep over multiple days.

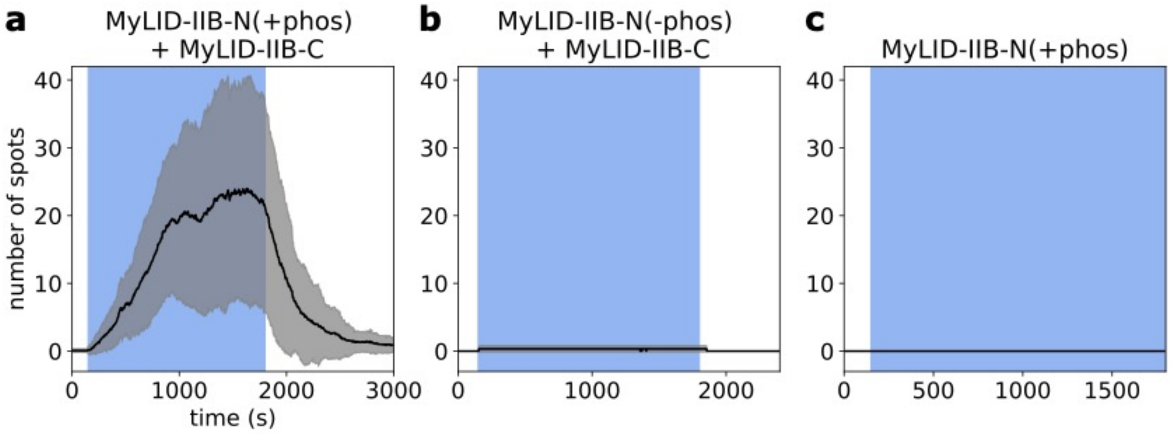

**Supplementary Figure 8. Control experiments for MyLID-IIB. a)** Number of moving minifilaments on the surface over time in a processive motility assay with phosphorylated MyLID-IIB-N and MyLID-IIB-C on biotinylated actin at 2 mM ATP in the presence of 0.3% methylcellulose (reproduced from Figure 4f). The black curve is the average of 13 movies across 13 different experiments, and the gray envelopes represent ± one standard deviation. **b)** Number of moving minifilaments on the surface over time in a processive motility assay with non-phosphorylated MyLID-IIB-N and MyLID-IIB-C under the same assay conditions as in **a)**. The black curve is the average of 3 movies across 3 different experiments. **c)** Number of moving minifilaments on the surface over time in a processive motility assay with only phosphorylated MyLID-IIB-N (MyLID-IIB-C was excluded from the assay) under the same assay conditions as in **a)**. The black curve is the average of 3 movies across 3 different experiments.

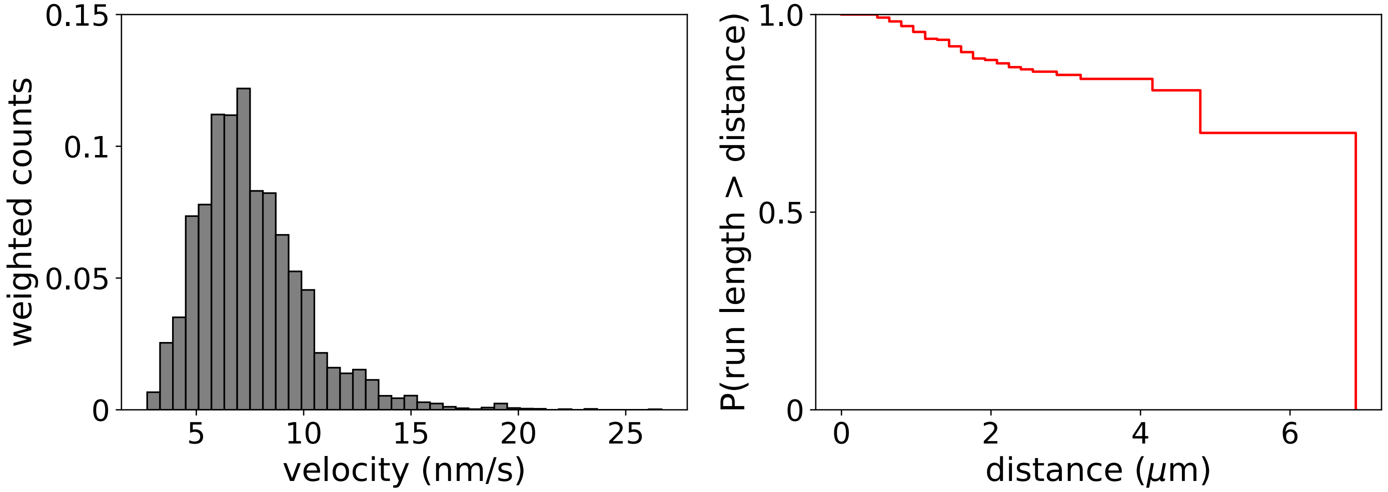

**Supplementary Figure 9. Velocities and run lengths for wildtype NMIIB minifilaments in processive motility assays on biotinylated actin.** Velocity histogram (left) and run length survival curve (right) for wildtype NMIIB minifilaments at 2 mM ATP in the presence of 0.3% methylcellulose. Data was collected from 13 separate experiments with 2 protein preps over multiple days.

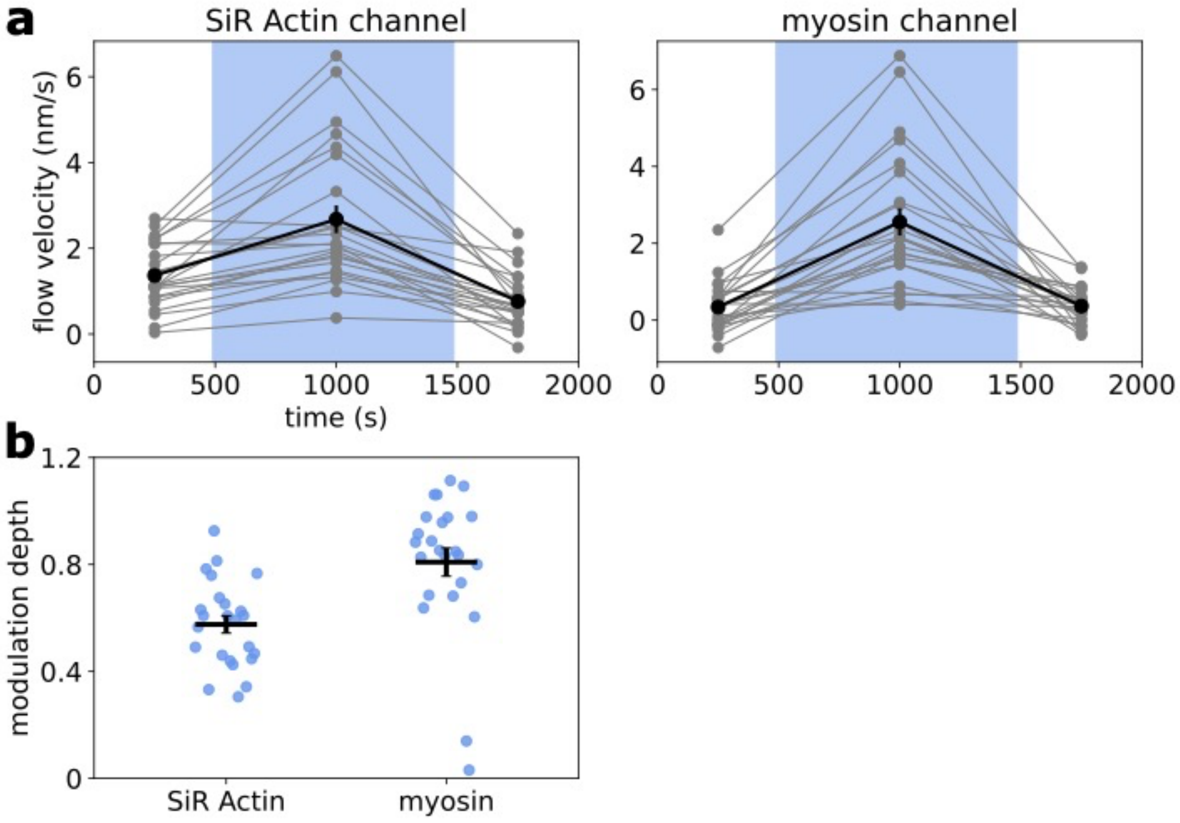

**Supplementary Figure 10. Flow velocity data plotted for individual cells. a)** Flow velocities for myosin (left) and SiR Actin (right) shown for each of the 24 cells that were used to calculate the mean flow rates reported in the text (same data set as Figure 5b). Each set of 3 connected gray points represents the mean flow velocities for a single cell before, during, and after blue illumination. Means were calculated by averaging over all frames within each of the 3 time periods of the blue illumination sequence (before illumination: 0-490 s, during illumination: 490-1490 s, and after illumination: 1490-1995 s). Black points represent the average across all cells within each time period. Error bars show standard errors. **b)** Modulation depth of flow velocities for each cell in the SiR Actin and myosin channels. Modulation depth is calculated as $1 - v_{dark}/v_{lit}$, where $v_{dark}$ is the time average over both the pre- and post-illumination periods, and $v_{lit}$ is the time average over the illumination period. Blue points represent modulation depths for each cell and black lines represent averages across all cells. Error bars show standard errors.

### Supplementary Movie Legends

**Supplementary Movie 1**
Processive motility assay of MyLOV-XI-IIB minifilaments on surface-immobilized biotinylated actin filaments at 500 µM ATP. The actin is shown in red and the myosin is shown in green. The static actin image is a maximum intensity projection of a short video of the actin that was acquired after imaging the myosin. Actin was labeled with Alexa 488 phalloidin and imaged with 488 nm excitation, and myosin was labeled with TMR Halotag ligand and imaged with 532 nm excitation. The blue background indicates when the blue LED was turned on.

**Supplementary Movie 2**
Processive motility assay of MyLOV-VI-IIB minifilaments on biotinylated actin filaments at 2 mM ATP. The actin is shown in red and the myosin is shown in green. The static actin image is a maximum intensity projection of a short video of the actin that was acquired after imaging the myosin. The white circles represent the plus-ends of actin filaments. Actin was labeled with Alexa 488 phalloidin and imaged with 488 nm excitation, and myosin was labeled with TMR Halotag ligand and imaged with 532 nm excitation. Actin polarity was determined by imaging unidirectional myosin VI tetramers labeled with Alexa 660 Halotag ligand using 633 nm excitation in the same field of view. The blue background indicates when the blue LED was turned on.

**Supplementary Movie 3**
Processive motility assay of MyLOV-VI-*zip* minifilaments on biotinylated actin filaments at 2 mM ATP. The actin is shown in red and the myosin is shown in green. The static actin image is a maximum intensity projection of a short video of the actin that was acquired after imaging the myosin. The white circles represent the plus-ends of actin filaments. Actin was labeled with Alexa 488 phalloidin and imaged with 488 nm excitation, and myosin was labeled with TMR Halotag ligand and imaged with 532 nm excitation. Actin polarity was determined by imaging unidirectional myosin VI tetramers labeled with Alexa 660 Halotag ligand using 633 nm excitation in the same field of view. The blue background indicates when the blue LED was turned on.

**Supplementary Movie 4**
Processive motility assay of MyLOV-$VI_{L310G}$-IIB minifilaments on surface-immobilized bundles of actin and fascin at 2 mM ATP. The myosin is shown in green, and the red channel shows a maximum intensity projection of the myosin as a representation of the actin bundles. The plus-ends of the actin bundles are oriented toward the top of the frame, and the minus-ends are oriented toward the bottom. Myosin was labeled with TMR Halotag ligand and imaged with 532 nm excitation. The blue background indicates when the blue LED was turned on.

**Supplementary Movie 5**
Contractility assay of MyLOV-XI-IIB minifilaments on a network of actin/alpha-actinin bundles at 2 mM ATP. The actin bundles were loosely attached to the surface via nonspecific interactions such that segments of the bundles remained unrestrained and could be deformed by the minifilaments. Only the actin channel is shown in this video. The myosin channel was imaged prior to the beginning of the video to confirm that minifilaments had bound to the actin bundles. Actin was labeled with Alexa 633 phalloidin and imaged with 633 nm excitation, and myosin was labeled with

TMR Halotag ligand and imaged with 532 nm excitation. The blue background indicates when the blue LED was turned on.

**Supplementary Movie 6**
Contractility assay of MyLOV-VI-IIB minifilaments on a network of actin/alpha-actinin bundles at 2 mM ATP. Only the actin channel is shown in this video. The myosin channel was imaged prior to the beginning of the video to confirm that minifilaments had bound to the actin bundles. Actin was labeled with Alexa 633 phalloidin and imaged with 633 nm excitation, and myosin was labeled with TMR Halotag ligand and imaged with 532 nm excitation. The blue background indicates when the blue LED was turned on.

**Supplementary Movie 7**
Processive motility assay of MyLID-XI minifilaments on biotinylated actin filaments at 10 µM ATP. The actin is shown in red and MyLID-XI-C is shown in green. MyLID-XI-N was not imaged in this video. The static actin image is a maximum intensity projection of a short video of the actin that was acquired before imaging the myosin. Actin was labeled with Alexa 488 phalloidin and imaged with 488 nm excitation. MyLID-XI-C was labeled with TMR Halotag ligand and imaged with 532 nm excitation, and MyLID-XI-N was labeled with Alexa 660 Halotag ligand. The blue background indicates when the blue LED was turned on.

**Supplementary Movie 8**
Processive motility assay of MyLID-IIB minifilaments on biotinylated actin filaments at 2 mM ATP. The actin is shown in red and MyLID-IIB-N is shown in green. MyLID-IIB-C was not imaged in this video. The static actin image is a maximum intensity projection of a short video of the actin that was acquired before imaging the myosin. Actin was labeled with Alexa 488 phalloidin and imaged with 488 nm excitation. MyLID-IIB-N was labeled with TMR Halotag ligand and imaged with 532 nm excitation, and MyLID-IIB-C was labeled with Alexa 660 Halotag ligand. The blue background indicates when the blue LED was turned on.

**Supplementary Movie 9**
Contractility assay of MyLID-XI minifilaments on a network of actin/alpha-actinin bundles at 10 µM ATP. The actin is shown in red and MyLID-XI-C is shown in green. MyLID-XI-N is not shown in this video. Actin was labeled with Alexa 633 phalloidin and imaged with 633 nm excitation. MyLID-XI-C was labeled with TMR Halotag ligand and imaged with 532 nm excitation, and MyLID-XI-N was labeled with Alexa 660 Halotag ligand. The blue background indicates when the blue LED was turned on.

**Supplementary Movie 10**
Imaging of a live S2 cell transfected with MyLOV-XI-IIB. The cytoplasm was labeled with CellTracker Deep Red to visualize cell morphology (shown in red) and MyLOV-XI-IIB was labeled with mRuby (shown in green). The cell was simultaneously imaged with 532 nm and 633 nm excitation. The blue background indicates when the blue LED was turned on. MyLOV-XI-IIB is localized to the lamella in this cell, and the lamella contracts toward the cell center during blue light exposure.

**Supplementary Movie 11**
Imaging of a live S2 cell transfected with MyLOV-XI-IIB. Actin was labeled with SiR Actin to visualize cell morphology (shown in red) and MyLOV-XI-IIB was labeled with mRuby (shown in green). The cell was simultaneously imaged with 532 nm and 633 nm excitation. The blue background indicates when the blue LED was turned on. MyLOV-XI-IIB is localized to the periphery of the lamella in this cell, and the lamella contracts toward the cell center during blue light exposure. Note that the untransfected cells in this field of view do not display blue light-dependent behavior.

**Supplementary Movie 12**
Imaging of a live S2 cell transfected with MyLOV-XI-IIB. The cytoplasm was labeled with CellTracker Deep Red (shown in red) and MyLOV-XI-IIB was labeled with mRuby (shown in green). The cell was simultaneously imaged with 532 nm and 633 nm excitation. The blue background indicates when the blue LED was turned on. MyLOV-XI-IIB is localized to thin protrusions in this cell, and the myosin flows toward the cell center during blue light exposure. Note that the protrusions cannot be seen in the CellTracker Deep Red channel due to poor labeling.

**Supplementary Movie 13**
Imaging of a live S2 cell transfected with MyLOV-XI-IIB. Mean flow maps for this cell are shown in Figure 5a (top row), and protrusions from this cell are shown in Figure 5d and Supplemental Video 15. Actin was labeled with SiR Actin to visualize cell morphology (shown in red) and MyLOV-XI-IIB was labeled with mRuby (shown in green). The cell was simultaneously imaged with 532 nm and 633 nm excitation. The blue background indicates when the blue LED was turned on. MyLOV-XI-IIB is mostly localized to protrusions and the lamella in this cell. The protrusions and their associated myosin retract inward toward the cell center during blue light exposure.

**Supplementary Movie 14**
Examples of contracting protrusions in live S2 cells transfected with MyLOV-XI-IIB. The left and right videos were taken from 2 separate cells during blue light exposure. The left video shows the protrusions depicted in the snapshots in Figure 5c. Actin was labeled with SiR Actin (shown in red) and MyLOV-XI-IIB was labeled with mRuby (shown in green). The cells were simultaneously imaged with 532 nm and 633 nm excitation. The protrusions shorten over time and there are occasional buckling events, indicating that contraction is occurring within the protrusions in these cells.

**Supplementary Movie 15**
Examples of contracting protrusions in live S2 cells transfected with MyLOV-XI-IIB. The left and right videos were taken from 2 separate cells during blue light exposure. The left video is taken from the cell shown in Supplementary Movie 13 and shows the protrusions depicted in the snapshots in Figure 5d. Actin was labeled with SiR Actin (shown in red) and MyLOV-XI-IIB was labeled with mRuby (shown in green). The cells were simultaneously imaged with 532 nm and 633 nm excitation. The lamella near the base of the protrusions contracts and the protrusions are pulled inward, indicating that lamellar contractility is partially responsible for retraction of protrusions in these cells.
